# Structural state recognition facilitates tip tracking of EB1 at growing microtubule ends in cells

**DOI:** 10.1101/636092

**Authors:** Taylor A. Reid, Courtney Coombes, Soumya Mukherjee, Rebecca R. Goldblum, Kyle White, Sneha Parmar, Mark McClellan, Marija Zanic, Naomi Courtemanche, Melissa K. Gardner

## Abstract

The microtubule binding protein EB1 specifically targets the growing ends of microtubules in cells, where EB1 facilitates the interactions of cellular proteins with microtubule plus-ends. Microtubule end targeting of EB1 has been attributed to high affinity binding of EB1 to GTP-tubulin that is present at growing microtubule ends. However, our 3D single-molecule diffusion simulations predicted a ∼6000% increase in EB1 arrivals to open, tapered microtubule tip structures relative to closed lattice conformations. Using quantitative fluorescence, single-molecule, and electron microscopy experiments, we found that the binding of EB1 onto opened, structurally disrupted microtubules was dramatically increased relative to closed, intact microtubules, regardless of hydrolysis state. Correspondingly, in cells, the conversion of growing microtubule ends from a tapered into a blunt configuration resulted in reduced EB1 targeting. Together, our results suggest that microtubule structural recognition, based on a fundamental diffusion-limited binding model, facilitates the tip tracking of EB1 at growing microtubule ends.

## Introduction

Microtubules are long, thin polymers that mechanically contribute to cell morphology, act as a track for molecular motor-based transport within the cell, and serve as a platform for binding of microtubule-associated proteins (Howard and Hyman, 2003; Mitchison and Kirschner, 1984; Ross et al., 2008). Microtubules are composed of αβ tubulin heterodimers stacked end-to-end into “protofilaments”. Typically, a microtubule is composed of thirteen laterally-associated protofilaments (Wang and Nogales, 2005; Zhang et al., 2015). Each tubulin heterodimer contains an exchangeable nucleotide site on the β-tubulin subunit, and individual tubulin heterodimers polymerize onto the plus-end of the microtubule with the β-tubulin subunit bound to a GTP nucleotide. After integration into the microtubule, the β-tubulin-bound nucleotide then stochastically undergoes hydrolysis and converts via GDP-Pi to GDP. GDP-bound tubulin is less stable in the microtubule lattice than GTP-bound tubulin, but the microtubule remains intact and continues to grow at its plus-end due to the continued addition of GTP-bound tubulin, which forms a “GTP cap” at the growing plus-end of the microtubule (Desai, 1997; Mitchison and Kirschner, 1984).

Recent literature suggests that microtubule structure may be more complex than previously considered, especially at the microtubule plus-end. A perfectly intact, closed microtubule lattice with “blunt” ends is defined by a regular arrangement of tubulin dimers into a thirteen protofilament tube, with all equal-length protofilaments terminating at the microtubule ends. In contrast, microtubule plus-ends have been observed by cryo-electron microscopy to have open, sheet-like or tapered conformations, which diverge greatly from a closed tube conformation (Chretien et al., 1995; Guesdon et al., 2016; Manka and Moores, 2018a). These findings have also been supported by quantitative analysis of fluorescence images (Coombes et al., 2013). This type of tapered, gently curved tip structure likely plays a role in the binding of Doublecortin and other proteins to microtubule plus-ends (Bechstedt and Brouhard, 2012; Bechstedt et al., 2014); reviewed in (Brouhard and Rice, 2014). Further, it has recently been reported that lattice damage and tubulin turnover can occur on the microtubule lattice itself, leading to irregularities along the length of a dynamic microtubule (Schaedel et al., 2015).

The microtubule tip tracking protein EB1 is a highly conserved protein that autonomously tracks the growing plus-ends of microtubules (Bieling et al., 2007; Dixit et al., 2009; Morrison et al., 1998). At the plus-end, EB1 recruits many other +TIP family proteins that have little to no native affinity for microtubules but that must localize to microtubule plus ends to perform their functions (Bieling et al., 2007; Dixit et al., 2009; Lansbergen and Akhmanova, 2006). High affinity binding of EB1 to the GTP-cap at growing microtubule plus-ends may contribute to its plus-end localization (Maurer et al., 2011; Zanic et al., 2009). The increased affinity of EB1 for GTP tubulin relative to GDP tubulin has been demonstrated through the use of GTP analogues, most commonly GMPCPP and GTP-γ-S. In both cases, there was an increase in overall EB1 binding to the GTP-analogue-bound microtubules as compared to GDP-microtubules (Maurer et al., 2011; Zanic et al., 2009). However, the mechanism for how EB1 rapidly and efficiently targets to growing microtubule plus-ends, thus allowing for robust tip tracking, remains unknown.

In this work we perfomed 3D single-molecule diffusion simulations, which predicted a ∼6000% increase in EB1 arrivals to open, tapered microtubule tip structures relative to closed lattice conformations. Using quantitative fluorescence, single-molecule, and electron microcopy experiments, we found that the binding of EB1 onto opened, structurally disrupted microtubules was dramatically increased relative to closed, intact microtubules, regardless of hydrolysis state. Further, we converted growing microtubule ends in LLC-Pk1 cells from a tapered to a blunt configuration, and observed a dose-dependent reduction in EB1 targeting. Together, our results suggest that microtubule structural recognition, based on a simple diffusion-limited binding model, facilitates EB1 tip tracking at growing microtubule plus-ends.

## Results

### At intermediate salt concentrations, EB1 preferentially binds to GMPCPP microtubule end structures

Previous in vitro studies have shown that while EB1 uniformly coats GTP-analogue (GMPCPP) microtubules in the absence of added salt, the addition of KCl increasingly drives EB1 off of the GMPCPP lattice (Dixit et al., 2009; Maurer et al., 2011; Zanic et al., 2009). Thus, we used an intermediate KCl concentration to explore the localization of EB1-GFP on reconstituted microtubules under relatively weak EB1 binding conditions (Fig. 1A, 30 mM KCl). We reasoned that under these conditions, any localized binding of EB1-GFP to GMPCPP microtubules could reveal an additional layer of regulation for EB1 binding, beyond exclusively tubulin subunit nucleotide state. Interestingly, we observed that, at 30 mM KCl, EB1 was often localized to GMPCPP microtubule ends (Fig. 1B, green; Fig. S1D).

**Figure 1:**
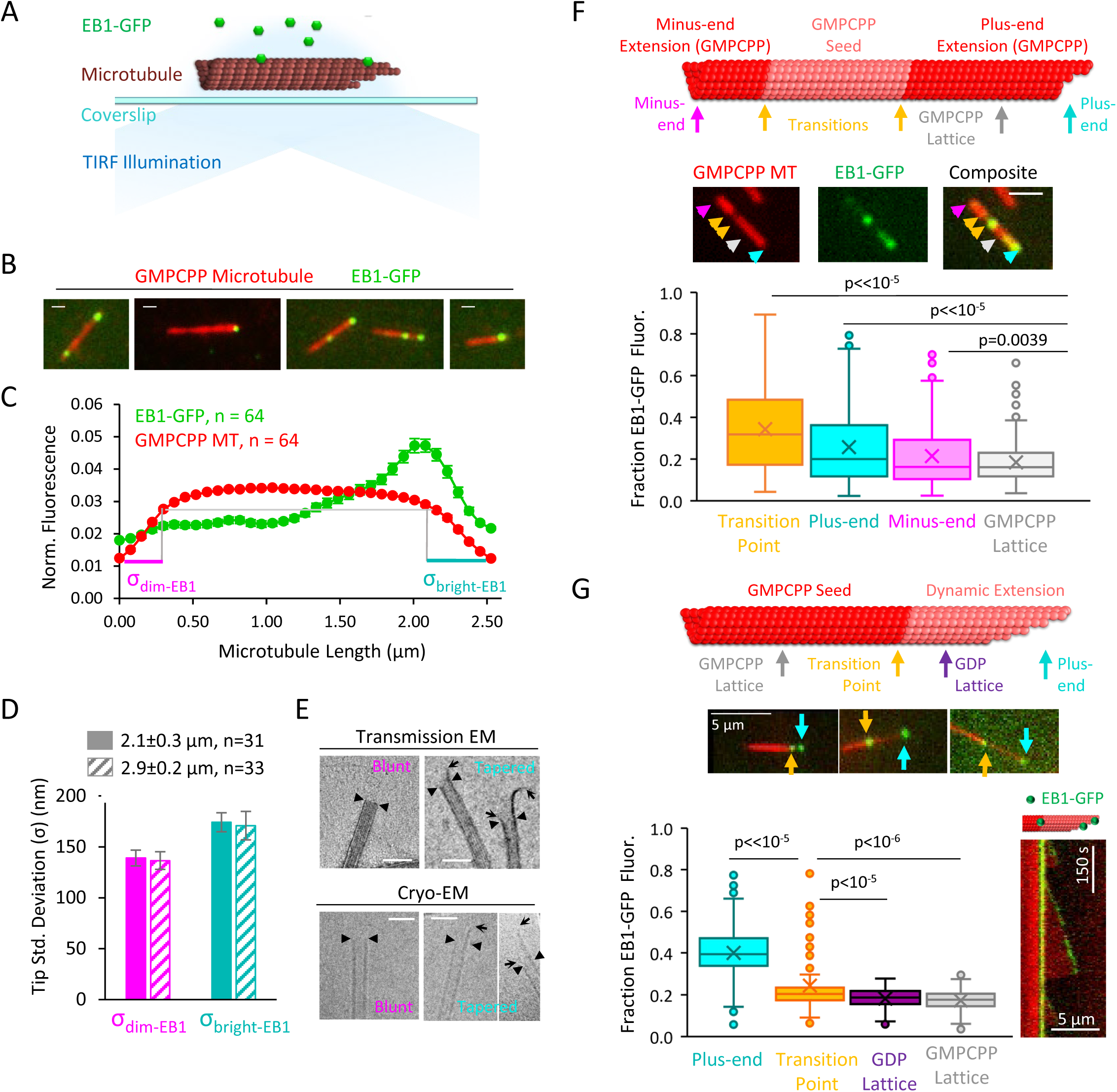
EB1-GFP recognizes structural disruptions and tapered ends on GMPCPP microtubules. (A) Schematic of EB1-GFP TIRF experiment. (B) Representative images of EB1-GFP (green) on rhodamine-labelled GMPCPP microtubules (red), showing preferential end localization. Scale bars: 0.4 μm. (C) Super-averaged intensity profile of 64 microtubules of length 2.5 ± 0.48 μm (mean±SD) with EB1-GFP. Individual microtubules within the averaged line scan were aligned with brighter EB1-GFP fluorescence on the right. Error bars show standard error at each point. (D) Bar graph with fitted tip standard deviations for microtubules of length 2.1 ± 0.3 μm (solid bars) and 2.9 ± 0.2 μm (hashed bars) (curve fits shown in Fig. S1). Lo ends (left), high tip standard deviation correspond to ends with more variable protofilament lengths (right). Error bars: SE calculated from 95% confidence intervals. (E) Electron microscopy images of blunt and tapered GMPCPP microtubules ends. Closed triangles indicate the most distal portion of the microtubule with a complete tubular lattice. Open arrows indicate the extrema of protruding protofilaments. Scale bars: 50 nm. (F) Top: Schematic of polarity-marked GMPCPP microtubule layout and reference positions for analysis. Center: Representative TIRF images of EB1-GFP on GMPCPP microtubules, colored arrows indicate the corresponding points used for analysis. Scale bars: 3 μm. Bottom: Box and whisker plot of the fraction of total EB1 fluorescence located at each position. The two transition points were averaged to provide a single value. Plus-ends, minus-ends, and transition points all showed significant differences as compared to lattice binding region (n=309 microtubules). (G) Top: Schematic of bright GMPCPP seed with dim dynamic microtubule extension, with reference positions for analysis. Center: Representative TIRF images of EB1-GFP on GMPCPP microtubules, colored arrows indicate the corresponding points used for analysis. Scale bars: 5 μm. Bottom-left: Box and whisker plot of the fraction of total EB1 fluorescence located at each position. Plus-ends and transition points both showed significant differences as compared to lattice binding region (n=134 microtubules). Bottom-right: Representative kymograph of EB1-GFP at transition point and plus-end during dynamic microtubule growth.

To detect whether there could be an underlying structure of the GMPCPP microtubule ends that would predispose them to EB1-GFP binding, we collected intensity line-scans of green EB1-GFP and red (rhodamine) GMPCPP microtubule fluorescence along the microtubules’ lengths to create average intensity profiles over all microtubules. The line-scans for 64 microtubules of similar length were rebinned to the mean microtubule length (see (Gardner et al., 2008) and Materials and Methods), and the ensemble average across all 64 microtubules was plotted for both the red and green channels (Fig. 1C). Importantly, prior to averaging, the line-scans in both the red and green channels for each microtubule were oriented such that the microtubule end with the brighter intensity of green EB1-GFP fluorescence was on the right side, with the dimmer EB1-GFP intensity on the left side (Fig. 1C, green, ensemble average over n=64 microtubules) (Coombes et al., 2016).

Similar to our qualitative observations, an increase in EB1-GFP intensity was observed at the brighter EB1-GFP microtubule end as compared to the center region and the dimmer EB1-GFP end of the microtubule (Fig. 1C). Strikingly, we noted that by aligning the brighter EB1-GFP intensity on the right side of the line scan, this resulted in a qualitative difference in the underlying microtubule intensity profile at the microtubule ends (Fig. 1C, red, ensemble average over n=64 microtubules). Here, the red microtubule fluorescence on the right (brighter EB1-GFP) side of the line-scan plot appeared to drop off more slowly to background as compared to the left side of the line-scan plot (Fig. 1C, compare teal vs magenta line lengths).

To quantify this observation, we fit a Gaussian error-function to the red fluorescence intensity drop-off at each microtubule end, as previously described (Coombes et al., 2013; Demchouk et al., 2011). This fitting process allowed us to estimate a “tip standard deviation”, which is a measure of tip tapering that arises as a result of protofilament length variability at the microtubule ends. To ensure that the tip fitting was not biased by rebinning a large range of microtubule lengths into one standard length, we subsampled two groups of microtubules (length standard deviation ≤ 0.3 μm in each case). Importantly, regardless of microtubule length, we found that the microtubule end with a brighter EB1-GFP signal had a larger average tip standard deviation, and thus more tip tapering due to protofilament length variation, than the opposite, dimmer EB1-GFP signal end, of the microtubule (Fig. 1D; 2.1±0.3 μm group (mean±SD): p=0.00017, Z-statistic = 3.79; 2.9±0.2 μm group (mean±SD): p=0.013, Z-statistic=2.47; Fit curves in Fig. S1A,B). We then used both Transmission Electron Microscopy (TEM) and Cryo-Electron Microscopy to verify that tapered and blunt ends were indeed present in the GMPCPP microtubule population (Fig. 1E), similar to previous observations (Atherton et al., 2018).

We then asked whether the inclusion of specific structural disruption points on the GMPCPP microtubules would alter EB1-GFP localization. Thus, polarity-labeled GMPCPP microtubule seeds were generated by growing bright-red GMPCPP extensions from dim-red GMPCPP seeds (Fig. 1F, top). These two-color microtubules allowed for identification of the microtubule plus-end as the end with the longer bright-red extension (Fig. 1F, top, teal). However, importantly, the polarity-labeled seeds also generated transition points between the original GMPCPP seeds and the GMPCPP extensions, which likely caused lattice discontinuities, defects, and holes at each transition point (Fig. 1F, top, yellow). Thus, we analyzed the brightness of EB1-GFP at 5 positions along the GMPCPP microtubules (Fig. 1F, top): the minus end tip (magenta), the two transition points (yellow), the plus-end tip (cyan), and the GMPCPP lattice (grey; for consistency and to minimize overlap with other positions, the lattice position used was always halfway between the plus-end and the nearest transition point). At each of the 5 positions, we summed the total green EB1-GFP intensity over a 9-pixel box, centered at the defined position, and then reported the fraction of EB1-GFP intensity at each position (= box intensity / summed intensity over all 5 boxes). For the transition point intensity, we reported an average of the two transition point box intensities for each microtubule. We found that the average transition point (p<<10^−5^, t-test), plus-end (p<10^−8^, t-test), and minus end (p=0.0039, t-test) of the GMPCPP microtubules all showed a significantly higher fraction of EB1-GFP fluorescence than the GMPCPP lattice position (Fig. 1G; n=309 microtubules).

Finally, to determine whether specific structural disruption points on the microtubule lattice could lead to EB1 binding even in the presence of dynamic microtubules, dim-red dynamic (GTP) microtubules were grown from bright-red GMPCPP seeds (Fig. 1G top) in the presence of 100 nM EB1-GFP and 55 mM KCl (Gell et al., 2010). We collected independent, individual microtubule images (Fig. 1G middle), and analyzed the brightness of EB1-GFP within a 9-pixel box at 4 positions along the microtubules (Fig. 1G, top and middle): the growing GTP-tubulin plus-end tip (cyan), the GDP lattice (purple), the transition point between the GMPCPP seed and the dynamic GDP microtubule extension (yellow), and the GMPCPP lattice (grey; note that for consistency and to minimize overlap with other positions, the GMPCPP and GDP lattice positions used were always halfway between the microtubule end and the nearest transition point). We found that, as would be expected, EB1-GFP intensity was ∼2.3-fold (230%) higher at growing microtubule plus-ends as compared to the overall average intensity on the GDP lattice and GMPCPP seeds (Fig. 1G bottom-left, cyan). However, consistent with our GMPCPP microtubule results, the EB1-GFP fluorescence at the GMPCPP/GDP transition point was 38% higher than the average intensity on the GDP lattice and GMPCPP seeds (Fig. 1G bottom-left; p<10^−5^ vs GDP lattice, p<10^−6^ vs GMPCPP seeds, n=134 microtubules). Further, in kymographs of dynamic microtubules, targeting of EB1-GFP to the GMPCPP/GDP transition occasionally persisted throughout entire microtubule growth and shortening events (Fig. 1G bottom-right).

### EB1 preferentially binds to disrupted-structure microtubules on GMPCPP, GTPγS, and GDP microtubule populations

Taken together, our GMPCPP and dynamic microtubule results suggested that EB1 may preferentially bind to regions of microtubules that have extended, tapered tip structures, or a discontinuous tubular lattice. To test this idea using microtubule populations with various tubulin-bound nucleotides, we used a previously published method to create pools of microtubules with common nucleotide states, but with varying degrees of lattice structural integrity (Reid et al., 2017). These protocols allowed for generation of “closed” microtubules with relatively intact, closed lattice structures (Fig. 2A, left), and “disrupted-structure” microtubules with lattice structures that had increased frequencies of defects, gaps in the lattice, and open sheet conformations (Fig. 2A, right, red subunits at structural disruptions). These “closed” and “disrupted-structure” populations were described for the three most commonly used in-vitro microtubule nucleotides (GDP, GMPPCP, and GTPγS (Fig. 2B)) (Reid et al., 2017), and so we used these populations as a tool to reveal potential differences in EB1 binding based on microtubule structure. Here, closed and disrupted-structure populations were made for GDP and GTPγS populations via taxol treatments, which created breaks, holes, and tapering in the microtubules, followed by differing storage conditions to alter the degree of overnight repair (Fig. 2B) (see (Reid et al., 2017) and Materials and Methods). In contrast, disrupted-structure GMPCPP microtubule populations were generated via Ca^2+^ treatment (Fig. 2B, center) (See (Reid et al., 2017) and Materials and Methods). Careful quantification that demonstrated a statistically significant structural disruption in the “disrupted-structure” populations as compared to the respective closed microtubule population in each case was previously performed using both fluorescent tubulin repair assays, and electron microscopy (Reid et al., 2017). We note that while the preparation protocols for microtubules using the three different nucleotides were distinct, each of these protocols reflected commonly used methods for producing stabilized in-vitro microtubules, and so no extreme or unusual conditions or preparation techniques were employed in preparing the microtubules.

**Figure 2:**
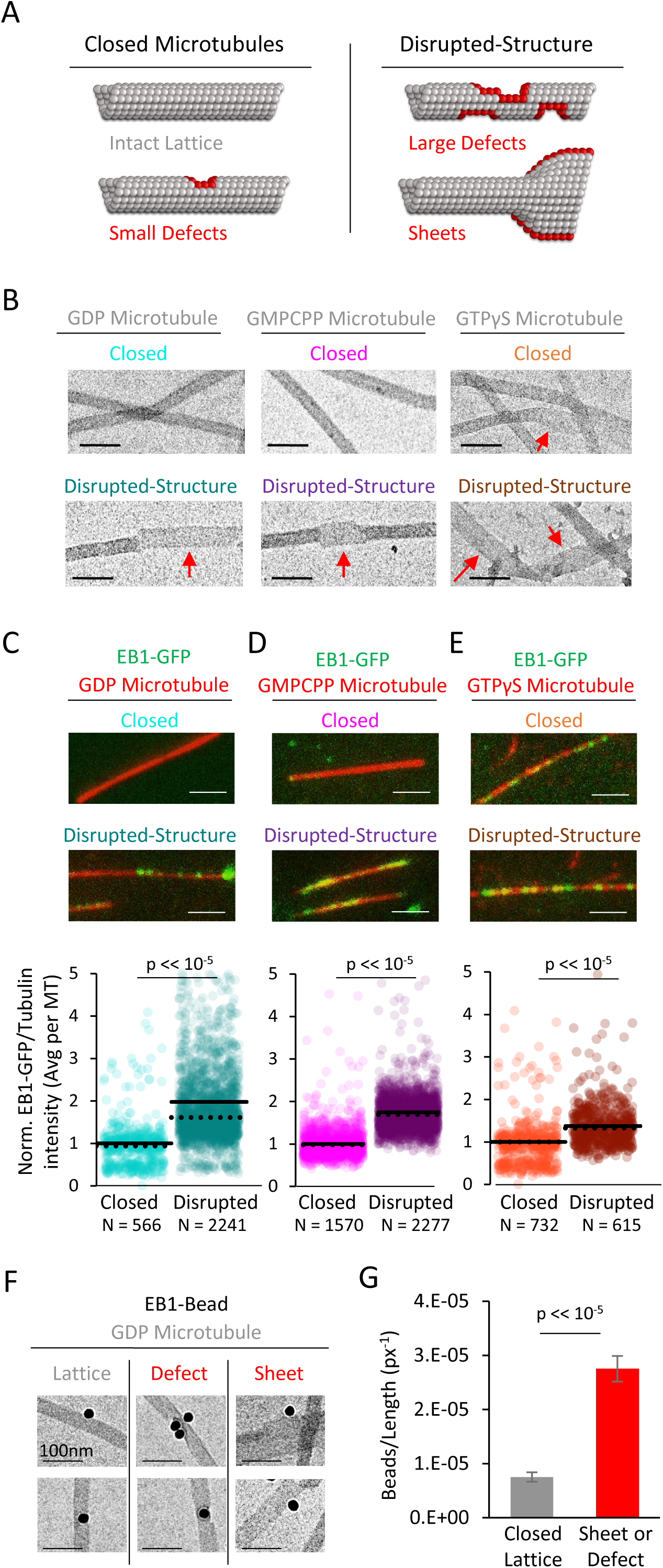
EB1-GFP shows preferential binding to disrupted-structure microtubules, regardless of nucleotide state. (A) Illustration of the two categories of microtubules used in this analysis. Left: “Closed” microtubules with predominantly intact, closed lattice structures. Right: “Disrupted-structure” microtubules have larger and more frequent gaps, defects, and sheet-like regions. Stylized red tubulin used to highlight defects. (B) Electron microscopy images of closed and disrupted-structure microtubules for GDP microtubules (left), GMPCPP microtubules (center), and GTPγS microtubules (right). Red arrows indicate structural disruptions, scale bars 50 nm. (C-E) Top: EB1-GFP (green) on microtubules (red) for closed and disrupted-structure conditions (scale bars 2 μm). Bottom: Cloud plots of normalized EB1-GFP/Tubulin binding ratio for individual microtubules, all values normalized to the grand average EB1-GFP/Tubulin binding ratio in the closed microtubule population. Each semi-transparent circle is the averaged data point from a single microtubule. Solid lines are the grand average EB1-GFP/Tubulin binding ratio for each population, dotted lines are the median EB1-GFP/Tubulin binding ratio for each population. (F) Electron microscopy of EB1 conjugated to gold beads on GDP microtubules. Example images of EB1-beads bound on the microtubule lattice (left), at a lattice defect (center), and on a sheet (right). (G) Frequency of EB1-beads bound to the microtubule lattice compared to EB1-beads bound at either a defect or on a sheet, normalized to observed microtubule length in each category. Non-normalized values yield a ∼1.7-fold increase in binding to defects and edges relative to a closed lattice (n=74 on lattice, n=128 on defect or sheet). Error bars SEM.

Thus, closed and disrupted-structure microtubule populations were generated for each nucleotide type, and were separately introduced into imaging chambers. Time was allowed for the microtubules to adhere to the coverslip surface, and then a solution of EB1-GFP and imaging buffer was introduced into the chambers (see Materials and Methods). After allowing time for EB1-GFP binding to reach steady-state (20-30 minutes), the microtubules with bound EB1-GFP were imaged using TIRF microscopy. We then reported the average EB1-GFP fluorescence intensity over background for each microtubule, which was then normalized to the average Rhodamine-tubulin fluorescence intensity over background for each microtubule. The average EB1-GFP/tubulin ratio for each individual microtubule was then normalized to the respective grand average EB1-GFP/tubulin ratio for the closed microtubule population in each case, to allow for direct comparisons between the disrupted-structure data and their respective closed microtubule population (grand average EB1-GFP/tubulin binding values for the closed microtubules populations (used for normalization) were as follows: GMPCPP: 0.098±0.0009; GDP: 0.100±0.006; GTPγS: 0.68±0.007 (mean±SEM); see Materials and Methods). We note that comparisons between different nucleotide types were not applicable in this experiment, due to dissimilar experimental protocols to generate microtubules for each nucleotide type (see Materials and Methods).

For the taxol-stabilized GDP microtubules, we observed a marked increase in the average ratio of EB1-GFP/tubulin bound to the disrupted-structure pool of microtubules as compared to the closed microtubules (Fig. 2C, top). Upon quantification, the average EB1-GFP/tubulin binding ratio in the disrupted-structure microtubule pool was ∼98% higher than the average EB1-GFP/tubulin binding ratio in the closed microtubule pool (Fig. 2C, bottom; p<<10^−5^, t-test). GMPCPP microtubules from the disrupted-structure pool also showed an increase in the average EB1-GFP/tubulin binding ratio (Fig. 2D, top), which was ∼75% higher than the average EB1-GFP/tubulin binding ratio in the closed microtubules (Fig. 2D, bottom; p<<10^−5^, t-test). Finally, the average EB1-GFP/tubulin binding ratio was higher on disrupted-structure GTPγS microtubules as compared to closed GTPγS microtubules, with a quantitative increase of ∼50% (Fig. 2E; p<<10^−5^, t-test).

### EB1 preferentially binds to disrupted-structure microtubules in electron microscopy experiments

To directly test whether EB1 preferentially binds to disrupted microtubule lattice structures on individual GDP microtubules, we used EB1 conjugated to gold beads, and examined the binding of these beads onto pre-stabilized microtubules using electron microscopy (Fig. 2F). For each microtubule-bound bead, the microtubule binding region was classified either as a complete lattice, or as a defect/sheet, and, further, the total available microtubule length for each classification was determined. Overall, we observed more EB1-gold beads that were bound to defects or sheets than to complete lattice regions (n=74 on lattice, n=128 on defect or sheet), regardless of total available length for each microtubule classification. However, when we normalized the frequency of bead location to the total available microtubule length for each classification, we found that the EB1-gold beads were nearly four-fold more likely to bind at defects or sheets than to a complete microtubule lattice (Fig. 2G; p<2.3×10^−54^, Chi-squared test).

### Single-molecule diffusion simulations predict a dramatic increase in EB1 on-rate to tapered microtubule structures

Taken together, our experimental results suggested that EB1 preferentially binds to regions of microtubules that have extended, tapered tip structures, or to a discontinuous tubular lattice. However, the mechanism for this preference remained unclear. Therefore, we developed a single-molecule diffusion-based computational simulation to explore a mechanistic explanation for our results.

For simplicity, we included three fundamental rules in the simulation: (1) EB1 molecules diffused in three dimensions with translational and rotational diffusion coefficients that depended on EB1 protein size (Fig. 3A, 1; see Materials and Methods) (Castle and Odde, 2013; Mirtich, 1998); (2) EB1 molecules could not pass through a microtubule, but rather would collide and then diffuse in another direction (Fig. 3A,2); and (3) the conformation of EB1, and its binding pocket within the microtubule lattice, were representative of their published molecular structures (Fig. 3A,3). To develop an approximation for the EB1 binding pocket within the microtubule, we used previously published Cryo-EM structural data as our guide (Fig. 3B, left; source data from (Zhang et al., 2015)), and from this data we created an approximation of the tubulin heterodimer shape (Fig. 3B, center), with careful consideration for EB1’s binding pocket, located between four tubulin dimers. EB1’s microtubule binding domain shape was similarly modeled from previously published Cryo-EM structures (Zhang et al., 2015), and the modeled shape was then verified to fit into our modeled 4-tubulin binding pocket in the correct orientation (3B, right). We note that each EB1 molecule could bind only one unique tubulin interface, and in only one unique orientation.

**Figure 3:**
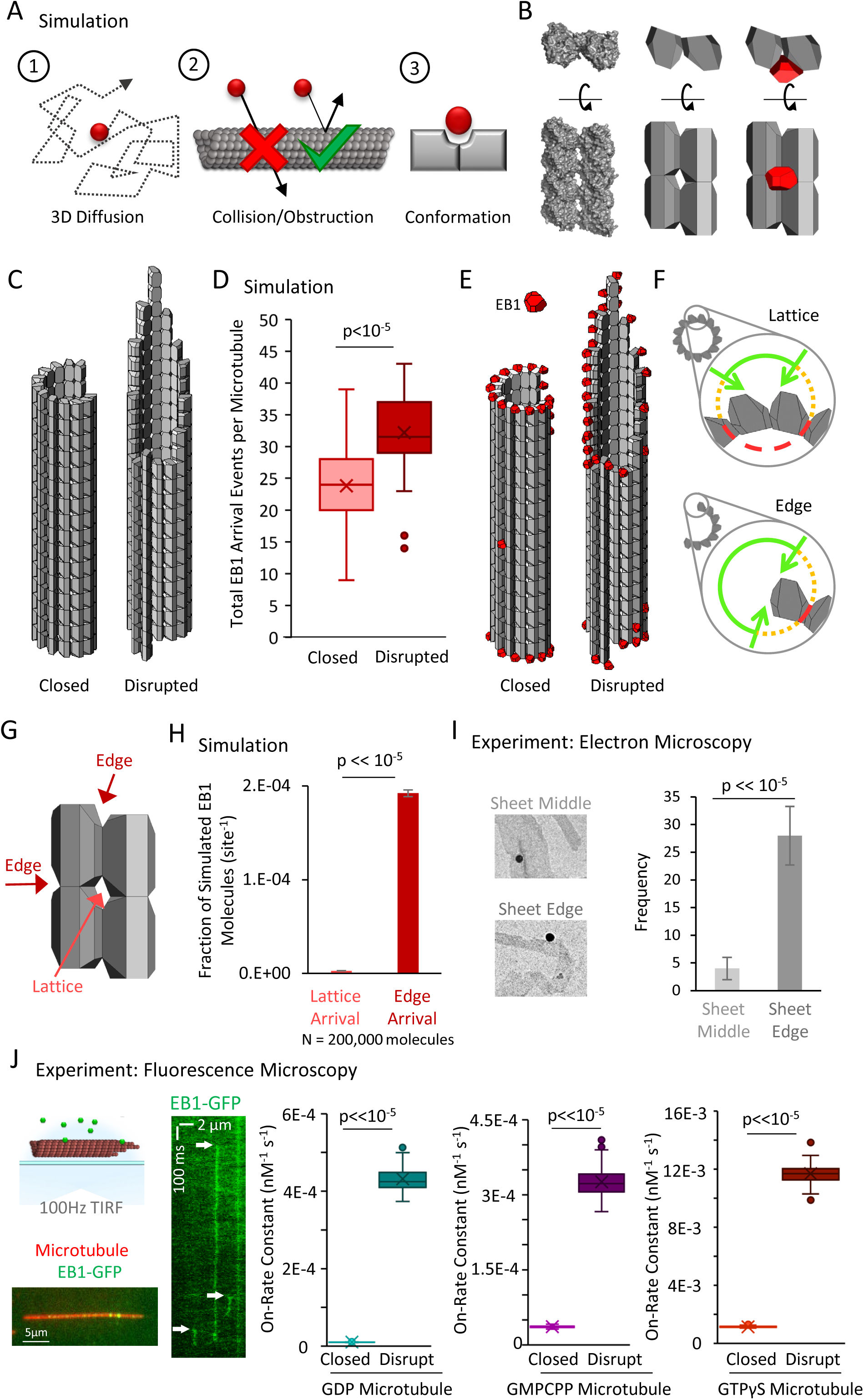
3D Diffusion-Based Simulations of EB1 predict an increased arrival rate at protofilament edge sites relative to closed, full lattice sites. (A) Simulation rules and parameters: (1) EB1 diffuses in three dimensions, (2) EB1 is obstructed by tubulin dimers and cannot pass through microtubules, (3) EB1 and tubulin dimer three-dimensional shapes reproduce EB1-tubulin binding interface. (B) Development of simulation shapes for EB1 and tubulin: Left: Cryo-EM reconstruction data (from PDB ID 3JAR) of tubulin dimers in a lattice: Left-top: Top-down view, with the lower portion being the outside of the microtubule. Left-bottom; Side view of four tubulin dimers, viewing the portion on the outside surface of the microtubule; Middle: Approximation of tubulin dimers for use in the simulation, derived from the cryo structure; Right: EB1 (red) also modeled as an approximation of cryo-structure data (not shown), with the binding interface correctly positioned at the pocket located between four adjacent tubulin dimers. (C) Two microtubule conditions used in the simulation analysis. Left: A closed, blunt-ended microtubule. Right: A “disrupted-structure” microtubule with a tapered tip. Both the “closed” and “disrupted structure” configurations contain the same total number of tubulin dimers. (D) Results of 50 simulations of 4,000 EB1 molecules each for each microtubule condition. Each data point in the box and whisker plot represents the total number of EB1 arrival events per microtubule for 4,000 different simulated EB1 molecules. (E) Visualization of EB1 arrival events (red) on closed and “disrupted-structure” microtubules from N=10,000 simulated EB1 in each condition. (F) Illustration of the hypothesis that EB1’s pocket-located binding site leads to a high local steric hindrance for EB1 binding to a lattice conformation (top), and a lower steric hindrance for EB1 binding to an edge conformation (bottom). Green lines: portion of the local volume with high accessibility to binding site. Yellow dotted lines: reduced accessibility volume. Red dashed lines: volume with no direct accessibility. In an edge conformation (bottom) the high accessibility region (green) is much larger than in the lattice conformation (top). (G) Dark Red arrows: Binding sites with two adjacent tubulin dimers, such as at the ends of protofilaments, or at exposed edges of protofilaments with no neighbor, are termed “Edge” sites. Light red arrow: Sites with four adjacent tubulin dimers are termed “Lattice” sites. Sites with only one tubulin dimer and with three adjacent tubulin dimers were not included in this analysis. (H) Fraction of simulated EB1 molecules that arrived at lattice sites (light red) or at edge sites (dark red). Data used was from the closed microtubule arrangement (panel C,E, left)). Values were determined by dividing the total number of EB1 bound at any lattice or edge site by the total number of available lattice or edge sites, respectively. EB1 was ∼70 fold more likely to bind a given edge site as compared to a given lattice site. (I) Left: example electron microscopy images of EB1 conjugated to gold beads on GDP microtubules. This image shows EB1-gold bound at a sheet edge, and one bound at a sheet middle (beads at ambiguous locations were conservatively classified as “middle”). Right: Count of total number of sheet bound beads observed over all images, divided into “Middle” of sheet and “Edge” of sheet. (J) Far Left Top; Experimental setup, rhodamine-labelled microtubules are affixed to a coverslip (red), EB1-GFP is introduced in solution (green), and the sample imaged at 100 frames per second using total internal reflection fluorescence (TIRF) microscopy. Far Left Bottom; Sample image of EB1-GFP on the microtubule. Left: Kymograph of EB1-GFP with length along the x-axis and time down the y-axis. White arrows indicate EB1-GFP binding events, which appear as vertical streaks. Example shows atypically long EB1 association events, for clarity. The lower limit of the vertical streaks are the dissociation event of EB1-GFP from the microtubule. Right: EB1-GFP on-rate constant for closed and disrupted-structure microtubules in each nucleotide population.

To run the simulation, a microtubule was fixed in space, and then individual EB1 molecules were allowed to diffuse, starting from random positions far away from the microtubule (see Materials and Methods). Two arrangements of microtubules, each with 207 tubulin dimers (average protofilament length ∼16 dimers), were simulated to mimic “closed” and “disrupted-structure” pools of microtubules. For the closed microtubules, a microtubule with a blunt end was defined (Fig. 3C, left), and for the “disrupted-structure” model, a microtubule with a tapered end was defined (Fig 3C, right). Starting from a random position away from the microtubule (see Materials and Methods), each EB1 molecule was then allowed to randomly diffuse either until it exceeded 2 μm away from the microtubule, or until the center of its binding interface was oriented properly and within 1 nm of each of its corresponding binding interface centers at any microtubule binding location (defined as an “arrival event”). Once either of these conditions was achieved for a particular EB1 molecule, the simulation for that molecule was ended, and, for EB1 molecules with arrival events, their arrival position on the microtubule was recorded. We note that off-rates (or dwell times) were not determined as part of this simulation, as a binding event concluded the simulation for an individual microtubule.

First, we ran simulations to evaluate the arrival events of EB1 molecules onto the two microtubule configurations, regardless of EB1-tubulin binding configuration (e.g., 1-4 tubulin binding interfaces were allowed per EB1 molecule). We ran 50 simulations of 4,000 EB1 molecules in each simulation for both the closed and the disrupted-structure microtubules, and calculated the total number of EB1 arrival events per microtubule in each simulation. We found that the mean number of successful arrival events of EB1 onto individual “disrupted-structure” microtubules was ∼ 33% higher than onto closed “blunt” microtubules with the same number of tubulin subunits (Fig. 3D; p<<10^−5^, Binomial test). Strikingly, by examining the simulated arrival locations of individual EB1 molecules, we found that the EB1 molecules had arrived at nearly all of the sites at the microtubule end or along laterally-exposed protofilaments, while very few arrived within a complete, closed lattice, both for the closed and the “disrupted-structure” microtubules (Fig. 3E, random representative subset of 10,000 simulated EB1 molecules).

Based on this observation, we hypothesized that the unique location of EB1’s binding site in the “pocket” between four tubulin dimers was responsible for the simulated “structural recognition” of EB1 (Fig. 3F, cutaway view – tubulin subunits above and below the section are not shown). Specifically, when a full lattice is present, the nature of the binding pocket between four tubulin dimers would restrict access of EB1 to its binding site, due to the high diffusional steric hindrance barrier generated by the nearby tubulin subunits, and the specific orientation that is required for EB1 to fit into this pocket and simultaneously bind at four locations (Fig. 3F, top, short green arc shows restricted access). However, when there is a region of the microtubule with exposed protofilament edges, the steric hindrance barrier would be dramatically reduced, thus allowing for increased accessibility and ease of EB1 binding to the laterally exposed edges (Fig. 3F, bottom, longer green arc shows increased access). This effect is likely even more dramatic than Fig 3F would suggest, since if this schematic were extrapolated to three dimensions, only a small wedge of a sphere would have high accessibility to an EB1 binding site for a closed lattice configuration. Importantly, a natural result of this model is that the on-rate of EB1 would be much higher at laterally (or longitudinally) exposed microtubule binding sites as compared to the binding pocket within a closed microtubule lattice.

To directly test the idea that the arrival rate of EB1 would be increased at laterally or longitudinally exposed microtubule binding sites, we ran simulations to compare the fraction of simulated EB1 arrival events onto closed-lattice microtubule binding sites with four adjacent tubulin dimers (Fig. 3G, light red arrow), to edge-located microtubule binding sites, with only two adjacent tubulin dimers either horizontally or vertically (Fig. 3G, dark red arrows). Thus, we specifically compared simulated EB1 arrivals to 2-tubulin vs 4-tubulin sites, excluding both1-tubulin sites that could potentially have a very high off-rate, and 3-tubulin sites. This allowed us to limit our analysis to the partial binding EB1 sites that have been previously observed using electron microscopy (Guesdon et al., 2016), and that we observed using electron microscopy in our new work (Fig. 2F).

Strikingly, in this simulation, the fraction of EB1 arrivals was ∼70-fold higher for 2-tubulin sites with lower steric hindrance, such as those at protofilament edges (Fig. 3H, dark red), as compared to 4-tubulin sites on the closed microtubule lattice (Fig. 3H, light red; p<<10^−5^, Chi-squared test). In order to compare these simulated arrival rates with the theoretical upper limit for our simulation, we modeled EB1 binding to a single, isolated tubulin heterodimer. In this instance, there would be no significant steric hindrance to binding, although arrival to the 1-tubulin site was still stereospecific, such that EB1 could not bind “upside down”. We found that the fraction of EB1 arrivals to these 1-tubulin sites was 309-fold higher than the fraction of arrivals to full lattice 4-tubulin sites, and 4.6-fold higher than the fraction of arrivals to 2-tubulin edge binding sites. This difference clearly demonstrates the large impact of steric hindrance for the pocket-located binding site of EB1 in the simulation, especially when fully enclosed within the lattice.

### Electron microscopy experiments demonstrate binding of EB1-gold beads to microtubule sheet edges

To test this model experimentally, we further analyzed the binding of EB1-beads onto stabilized microtubule sheets from our electron microscopy experiments (Fig. 3I, left). Qualitatively, we observed EB1-beads that bound to the stabilized microtubule sheets, both in the middle of the sheet, and on the edges of sheets (Fig. 3I, left), as has been previously observed on the more transient microtubule sheets that are present at dynamic microtubule plus-ends (Guesdon et al., 2016). To quantify this observation, we counted the number of beads located at sheet edges as compared to the middle of sheets. Here, beads that were ambiguously located were conservatively classified as being at the middle of the sheet. We observed that, for sheet-like regions of microtubules, there were 7-fold more beads associated with sheet edges than with the middle of the sheet (Fig. 3I, right; p<<10^−5^, Chi-squared test). This suggests that EB1 can indeed bind at edge sites on a microtubule, despite the reduced number of inter-site binding partners. Further, the ratio of edge sites to total binding sites within a sheet is estimated at 2:14 (edge sites to total binding sites), and so by normalizing to the number of available sites in each case, this yields an observed ∼49-fold preference for edge sites over lattice sites. This fold preference is on the order of the ∼70-fold increase in EB1 on-rates for edge sites as compared to lattice sites from the 3D diffusion simulation (Fig 3H). We note that there was a lower reported ratio of edge to middle position binding on dynamic microtubule sheets (1:18, (Guesdon et al., 2016)). However, for growing microtubules in the presence of free tubulin, the possibility that EB1 first binds to a sheet edge, and that this edge binding position is then converted to a sheet-middle position by subsequent addition of new tubulin subunits, cannot be ruled out. In contrast, for our stabilized microtubules with no free tubulin, binding in a sheet-middle position would require that EB1 binds directly to the sheet-middle position. Therefore, the mechanism for localization of EB1 to sheet-middle positions may be very different for stable microtubule experiments with no free tubulin, as compared to dynamic, growing microtubule experiments in the presence of free tubulin.

### Rapid frame-rate fluorescence microscopy experiments demonstrate increased on-rate of EB1-GFP to disrupted-structure microtubule pools

Our 3D single-molecule diffusion simulations predicted a ∼70-fold rise in the fraction of EB1 arrivals to 2-tubulin edge structures relative to a 4-tubulin-pocket closed lattice conformation, due to a high diffusional steric hindrance barrier that frustrates EB1 from binding in the pocket-like interface between four adjacent tubulin dimers in the lattice. Thus, the simulated arrival rate of EB1 to its binding site on the microtubule was dictated by the structural state of the microtubule itself. To experimentally test this model prediction, we collected rapid frame-rate (100 Hz) movies using TIRF microscopy, and then tracked the association and dissociation events of individual EB1-GFP molecules with microtubules (Fig. 3J, left; white arrows in EB1-GFP kymograph indicate EB1 association events). Because it was not possible to directly distinguish between EB1-GFP that was bound to a protofilament edge vs EB1-GFP that was bound at a closed lattice site in our TIRF experiments, we used our closed and disrupted-structure microtubule preparation protocols (Fig. 2) to measure the difference in EB1-GFP association and dissociation rates between closed microtubules, and microtubules that likely had an increased number of exposed edge binding sites (disrupted-structure microtubules). The EB1-GFP on-rate was calculated in each case by fitting a decaying exponential to the dwell time histogram for individual binding events (Fig. S2), which allowed us to calculate the expected total number of binding events, including those with a very short dwell time (Telley et al., 2009) (See Materials and Methods).

We found that the association rate for EB1-GFP was ∼45-fold higher for disrupted-structure microtubules as compared to closed microtubules for the GDP microtubule pools (Fig. 3J, center, teal; p<<10^−5^; Z-test; normalized data Fig. S2E). This fold increase was similar to the normalized results for the electron microscopy EB1-bead studies (Fig. 3I), suggesting that there was a substantial increase in edge-binding sites in the disrupted-structure pool as compared to the closed pool. For the GMPCPP microtubule pool, there was a ∼12-fold increase in the EB1-GFP association rate constant in the disrupted-structure microtubule pool relative to the closed pool (Fig. 3J, center, purple; p<10^−64^; Z-test; normalized data Fig. S2F), suggesting that the CaCl_2_ treatments used to disrupt microtubule structure may be less efficient at producing edge-binding sites than the protocol for the GDP microtubule pools, which relied on damage initiated by Taxol treatment. Finally, the EB1-GFP association rate constant was ∼3.5-fold higher in the disrupted-structure microtubule pool as compared to the closed microtubules for the GTPγS microtubules (Fig. 3J, right, brown; p=1.7×10^−16^; Z-test; normalized data Fig. S2G). This more moderate increase in the association rate constant may be due to a smaller difference in microtubule structure between the closed and disrupted-structure pools of GTPγS microtubules, as the closed microtubule pool itself was shown to exhibit structural disruption (Reid et al., 2017).

We then calculated EB1-GFP dwell times on the microtubule lattice from fits to our experimentally observed exponentially decaying dwell-time histograms (Fig S2A). We found that all pools of disrupted-structure microtubules had lower characteristic dwell-times than their respective closed pools (Fig S2B), suggesting that, while EB1 binding to protofilament edges may be rapid due to low steric hindrance, a 2-fold reduction in the number of EB1-tubulin bonds at the edge binding sites would lead to an increased EB1 off-rate. Consistent with this model, we found that EB1 dwell time distributions were best modeled as two exponential distributions (Fig S2C,D), one with a short dwell time (∼20 ms), which could be associated with edge-bound EB1, and a second underlying distribution with a longer dwell time (∼150 ms), which may correspond to lattice-bound EB1.

### In experiments and simulations, a tubulin face-binding protein does not exhibit microtubule structure recognition

Taken together, our data suggests that the pocket localization of EB1’s binding site on the microtubule acts to limit access of EB1 to 4-tubulin lattice binding sites relative to 2-tubulin edge binding sites (Fig. 4A, left vs right, lighter green arc shows large increase in accessibility). In contrast, this model predicts that a protein with a binding site on the face of a tubulin dimer would have similar binding site accessibility on both lattice and edge sites (Fig. 4B, left vs right, lighter green arc shows small difference in accessibility). Thus, we predicted that a tubulin face binding protein would not show an increased number of arrival events to disrupted-structure microtubules.

**Figure 4:**
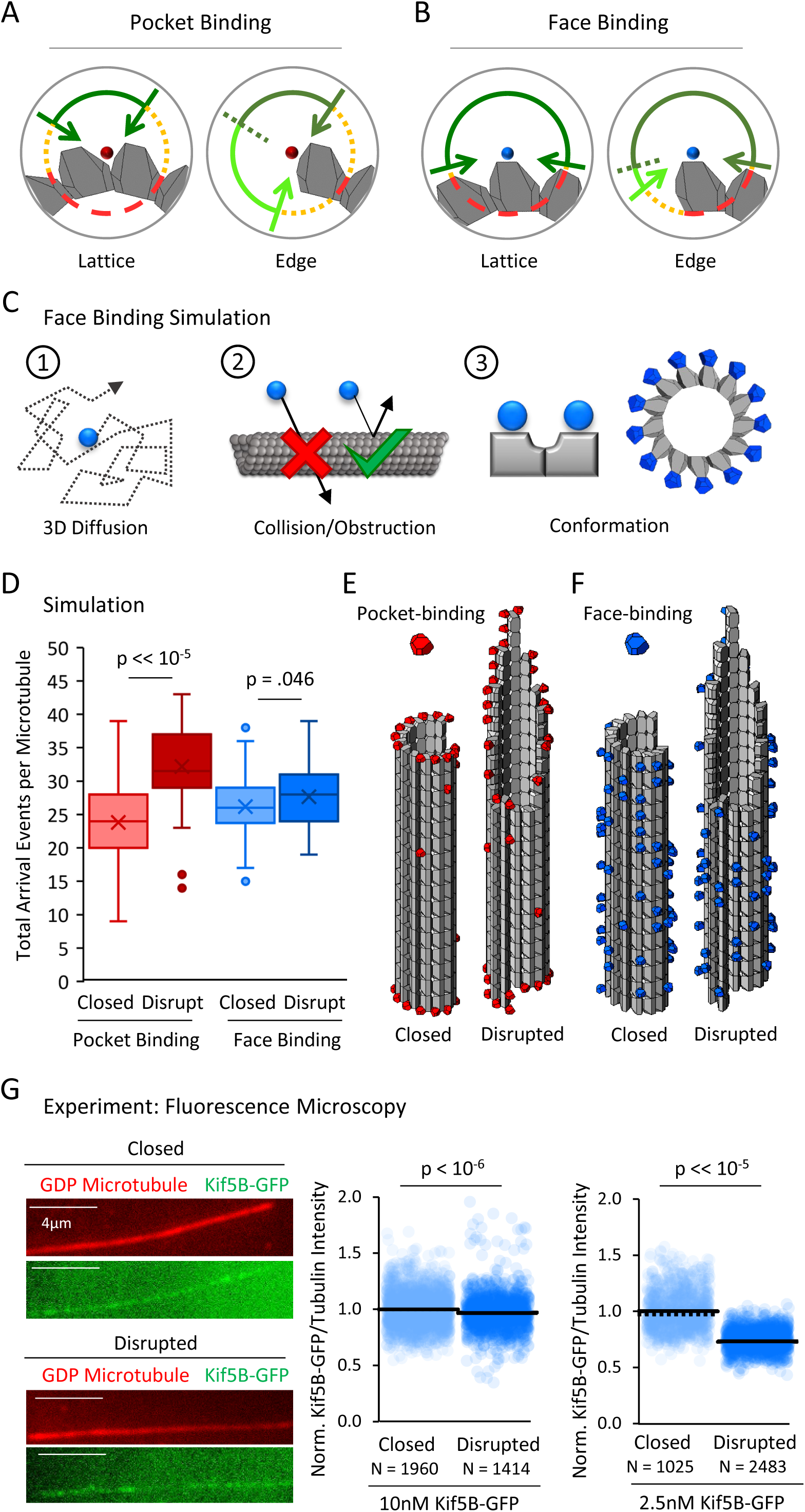
A tubulin face binding protein does not demonstrate microtubule structure recognition in experiment and simulation. (A) Hypothesis schematic showing large increase in direct access to a pocket binding site (dark green + light green) upon change from lattice conformation (left) to an edge conformation (right) for a pocket-binding protein. (B) Face binding hypothesis model predicts only a small change in direct access to a face binding site upon change from lattice conformation (left) to an edge conformation (right). (C) Simulation rules and parameters for face binding are identical to a pocket binding simulation with respect to (1) 3D diffusion, and (2) obstruction of the binding molecule by tubulin molecules. (3) The only change in the simulation was in the location of the binding site (left), with the arrangement of simulated face-binding protein on a microtubule at right. (D) Total arrival events per microtubule for a simulated pocket binding protein (EB1, red) and face binding protein (blue), showing molecular arrival events on closed (light red / light blue) and “disrupted-structure” (dark red / dark blue) microtubule conformations. Results of 50 simulations of 4,000 EB1 molecules each for each microtubule condition. Each data point in the box and whisker plots represents the total number of EB1 arrival events per microtubule for 4,000 different simulated EB1 molecules. (E) Representative visualizations of arrival event location distribution of pocket binding protein such as EB1 (red) on closed and disrupted-structure microtubules. (F) Representative visualizations of arrival event location distribution of face binding protein (blue) on closed and disrupted-structure microtubules, illustrating the loss of preference for edge-located binding sites. (G) Left: Representative images of Kif5B-GFP (green), which is a tubulin face binding protein, on GDP microtubules (red) (scale bar 4 µm). Right: Cloud plots of normalized Kif5B-GFP/Tubulin binding ratio for individual microtubules, all values normalized to the grand average Kif5B-GFP/Tubulin binding ratio in the closed GDP microtubule population. Each semi-transparent circle is the averaged data point from a single microtubule. Solid lines are the grand average Kif5B-GFP/Tubulin binding ratio for each population, dotted lines are the median EB1-GFP/Tubulin binding ratio for each population.

To ask whether our simulation supported this prediction, we first performed tubulin “face binding” simulations that had identical rules to the EB1 diffusion simulations (Fig. 4C, 1-2), but with the exception that the protein binding site was located on the outer face of a tubulin dimer (Fig. 4C, 3), rather than inside the four-tubulin pocket between dimers. Thus, we used our previous tubulin dimer and EB1 conformations, but instead moved the protein binding location to the outer face of the tubulin heterodimer, instead of within the 4-dimer pocket between tubulin dimers (Fig. 4C, 3, right). We then asked how the total number of face-binding molecule arrival events per microtubule in the closed and disrupted-structure microtubule simulations would be altered by changing this rule.

In previous EB1 simulations, the mean number of EB1 arrival events per microtubule was increased for the disrupted-structure microtubules relative to the closed microtubules (Fig. 4D, red; p=3.2×10^−28^, Binomial test; 4000 EB1 molecules per simulation, 50 simulations). In contrast, we found that when the protein binding site was located on the tubulin heterodimer’s outward face, the mean number of face-binding molecule arrival events for disrupted-structure microtubules was not substantially increased relative to the closed microtubules (Fig. 4D, blue; p=.046, Binomial test; 4000 face-binding molecules per simulation, 50 simulations), consistent with our hypothesis that the face-binding protein would not differentiate between edge and lattice binding sites. Similarly, the previous observation that simulated, pocket-binding EB1 molecules showed preferential arrivals to edge-proximal tubulin heterodimers (Fig. 4E, red) was strikingly altered in the tubulin face-binding simulations: the face-binding protein arrival events were located non-specifically along the lattice, and with no enrichment on edge dimers (Fig. 4F, blue; random representative subset of 10,000 simulated EB1 molecules). Thus, our simulations predicted that a tubulin face-binding protein would show similar rates of arrival to both closed and disrupted-structure microtubule populations, due to a similar steric hindrance to binding in both cases.

To experimentally test this prediction, we used a monomeric Kif5B-GFP kinesin construct in the presence of 10 mM AMPPNP, to allow for microtubule binding analysis of the monomeric kinesin (K339-GFP) (Case et al., 2000; Tomishige and Vale, 2000). The Kinesin-1 Kif5B protein has been shown to dock with a stoichiometry of one kinesin motor per tubulin heterodimer, and binds to the outer surface of the α and β tubulin monomers, rather than the lateral surfaces, in electron microscopy experiments (Hirose and Amos, 1999; Kikkawa et al., 2000; Kozielski et al., 1998; Lowe et al., 2001; Marx et al., 2006; Moores et al., 2002; Nogales et al., 1999; Rice et al., 1999; Sosa et al., 1997). Thus, we used the Kif5B-GFP construct to determine whether a tubulin face-binding protein would show differential binding to closed and disrupted-structure microtubule populations, as was observed for the EB1-GFP protein (Fig. 2B).

We first used 10 nM Kif5B-GFP, and compared overall binding of the Kif5B-GFP to closed and disrupted-structure pools of GDP microtubules, similar to the procedure for our EB1-GFP experiments (Fig. 2B). Qualitatively, and in contrast to the EB1-GFP experiments, the degree of binding between the closed and disrupted-structure pools of GDP microtubules appeared similar (Fig. 4G, left). Quantification of the TIRF images demonstrated that the average Kif5B-GFP/tubulin binding ratio for the disrupted-structure pool of GDP microtubules was not higher than in the closed microtubules (p=1; one-sided t-test), but rather appeared to be slightly lower than in the closed microtubules (p = 2.7×10^−7^, 2-sided t-test), perhaps suggesting a slight suppression of Kif5B-GFP binding to microtubules with disrupted-structure structures. Similar results were observed using a lower concentration of Kif5B-GFP (2.5 nM, Fig. 4G, right; p<10^−15^, two-sided t-test). These results suggest that the binding location of EB1 on microtubules may dictate its preferential binding to the disrupted-structure pool of microtubules relative to the closed microtubules, and that binding to the face of a tubulin dimer by Kif5B does not yield this preferential binding behavior.

### EB1 tip tracking by microtubule structure recognition

We then asked how the rapid binding of EB1 onto protofilament edges could be integrated with current models for tip tracking behavior on growing microtubule ends. It has been previously shown that EB1-GFP is concentrated in a location on growing microtubule ends that is slightly behind the distal tip (Maurer et al., 2014). This EB1-GFP fluorescence distribution has been explained via a model in which EB1 first binds the microtubule with high affinity in a location that is penultimate to the microtubule tip, and then, over time, a conformational change occurs on the microtubule that lowers the affinity of EB1 for this binding site (Maurer et al., 2014). Because EB1 binds the microtubule with high affinity in a location that is penultimate to the microtubule tip, one hypothesis is that EB1 binds with high affinity to a GDP-P_i_ lattice, which would be localized penultimate to the tip of the microtubule due to a delay in hydrolysis of newly added GTP-tubulin subunits at the microtubule tip (Zhang et al., 2015). In this model, the conversion of the microtubule lattice from a GDP-P_i_ state to a GDP state would then lower the affinity of EB1 for its binding site. Thus, we analyzed the overall binding of EB1-GFP to GDP-P_i_ microtubules as compared to GDP microtubules, to determine whether EB1-GFP binds with high affinity to GDP-P_i_ microtubules, and with a lower affinity to GDP microtubules.

It has been previously shown that the K_d_ of P_i_ for GDP is ∼25 mM (Carlier et al., 1988; Raw et al., 1997). Furthermore, the depolymerization rate of GDP microtubules is substantially slowed in the presence of 50 mM of inorganic phosphate, indicating that P_i_ binds to GDP-tubulin under these conditions (Carlier et al., 1988). Therefore, under these conditions, we compared the binding of EB1-GFP to GDP and GDP-P_i_ microtubules to determine whether EB1 exhibits preferential binding to GDP-P_i_ microtubules (Fig. 5A; see Materials and Methods). Surprisingly, EB1-GFP binding appeared to be reduced on GDP-P_i_ microtubules relative to GDP microtubules, for both closed and disrupted-structure microtubules (Fig. 5B). By quantifying EB1-GFP binding on each microtubule type using our previously described automated analysis tool (Reid et al., 2017), we found that there was a −53% reduction in binding of EB1-GFP to closed GDP-P_i_ microtubules relative to closed GDP microtubules (p<<10^−5^, t-test) (Fig. 5C, experimental repeats Fig. S3A). These results could perhaps be explained by previous cryo-electron microscopy findings in which it was demonstrated that the GDP-P_i_ microtubule structural state is distinct from that of GDP microtubules (Manka and Moores, 2018b). We conclude that EB1 does not directly bind to GDP-P_i_ microtubules, but, instead, that hydrolysis of GTP-tubulin to GDP-P_i_-tubulin could initiate formation of a low-affinity EB1 binding site.

**Figure 5:**
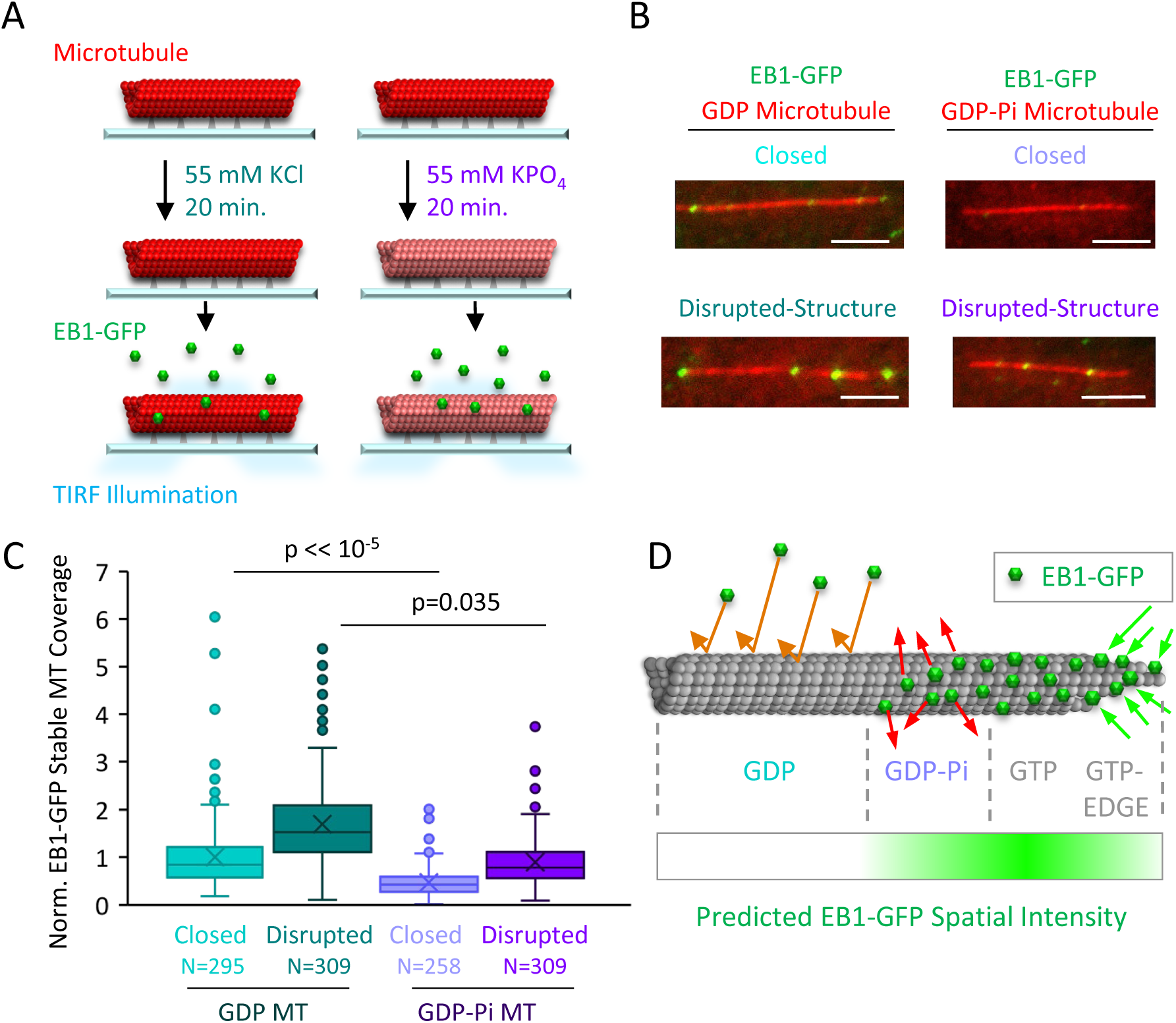
EB1-GFP binding is suppressed on GDP-P_i_ microtubules. (A) Experiment to compare EB1-GFP binding to GDP and GDP-P_i_ microtubules (see Materials and Methods). (B) Representative images of EB1-GFP binding to microtubules (scale bar 5 µm). (C) Quantitative binding comparison, based on EB1-GFP binding area on microtubules. All values normalized to the grand average binding area in the closed GDP microtubule population. Sample sizes represent number of independent images analyzed. (D) Implementation of structural recognition into a 2-state model for EB1 binding and microtubule tip structure maturation.

Our finding that EB1 binds with low affinity to GDP-P_i_ lattices is perhaps unexpected based on previous dynamic microtubule experiments in which it was observed that Mal3-GFP was bound to the entire dynamic microtubule lattice in the presence of GTP together with BeF_3_^−^ (Maurer et al., 2011). Here, while BeF_3_^−^ is thought to replace the dissociated phosphate on the GDP lattice, perhaps mimicking a GDP-P_i_ state, it may be that BeF_3_^−^ instead acts to lock the dynamic microtubule lattice into a GTP-like state in the presence of excess GTP, which is required in a dynamic microtubule assay. However, GTP was absent in our GDP-P_i_ microtubule experiments, thus ensuring that phosphate was bound only to the GDP lattice.

Thus, our GDP-P_i_ data, combined with our protofilament edge binding data, led us to hypothesize that the process of EB1 binding at the growing microtubule end could be characterized by (1) rapid EB1 binding to edge sites at the distal end of the microtubule, followed by maturation into a tight EB1 binding configuration through the addition of new GTP-tubulin subunits at locations of edge-bound EB1, which would stabilize EB1 binding by completing its 4-tubulin pocket binding configuration (Fig. 5D, right), and (2) subsequent destruction of the stable 4-GTP-tubulin pocket binding site by hydrolysis of GTP-tubulin into GDP-P_i_ (Fig. 5D, center), and, ultimately, GDP-tubulin, which may be accelerated by the binding of EB1 into its 4-tubulin pocket binding configuration (Maurer et al., 2014; Zhang et al., 2015; Zhang et al., 2018).

We predict that rapid binding to protofilament edge sites at tapered microtubule ends would remain consistent with observations that the highest density of EB1-GFP is 80-160 nm behind the most distal end of the microtubule (Maurer et al., 2014). Here, although new edge sites are continuously generated as a result of tubulin subunit additions to growing microtubule plus-ends, the density of edge sites on an open, tapered microtubule sheet is, by definition, 7-fold lower than that of lattice sites on a closed microtubule tube (ratio of edge-binding-sites to total-binding-sites per tubulin layer for an open microtubule sheet is 2:14). Thus, even with rapid loading of EB1 onto edge sites at the most distal positions on the microtubule tip, the peak in EB1-GFP intensity for a plus-end EB1-GFP comet would be penultimate to the tapered tip, in the location where a high density of EB1 molecules are bound to the much higher density, four-GTP-tubulin pocket lattice binding sites, perhaps via addition of new, incoming GTP-tubulin subunits onto the location of previously edge-bound EB1 molecules. Consistent with this prediction, by generating simulated microscope images of microtubule tips with complete occupancy of EB1-GFP at distal tip edge sites and an exponentially decaying occupancy of EB1-GFP away from the distal tip on the closed microtubule lattice sites (Fig. S3B), we observed that the highest density of EB1-GFP lagged behind the distal microtubule end (Fig. S3C), as previously described (Maurer et al., 2014). Thus, in a model where EB1 is preferentially targeted to edge binding sites at tapered microtubule ends, the reduced density of edge binding sites relative to closed microtubule lattice binding sites could explain the penultimate location of EB1 at growing microtubule ends (Fig. 5D).

### Flattening of tapered microtubule tip structures suppresses EB1 binding in cells

An important prediction of a structural-recognition model for EB1 trip tracking at growing microtubule ends (Fig. 5D) is that the pruning of extended, tapered tip structures at growing microtubule ends, leading to blunt ends with fewer protofilament-edge EB1 binding sites, would suppress EB1 tip tracking in cells. To test this idea, we treated LLC-Pk1 cells that expressed Tubulin-GFP (Rusan et al., 2001) with increasing, low dose concentrations of the microtubule destabilizing drug Vinblastine, to determine whether we could alter the structure of the growing microtubule plus-ends. While Vinblastine does not alter the hydrolysis rate of GTP-tubulin (Castle et al., 2017), it interferes with tubulin assembly into protofilaments (Gigant et al., 2005), and could thus suppress the tapered multi-protofilament extensions that have been observed at rapidly growing microtubule plus-ends (Chretien et al., 1995; Coombes et al., 2013; Guesdon et al., 2016).

We treated LLC-Pk1 cells with low concentrations of Vinblastine, which still allowed for robust growth of microtubules (Fig. 6A). Then, we collected line scans of the drop-off in tubulin-GFP fluorescence at growing microtubule plus-ends, to assess whether microtubule tip structures were altered (Fig. 6B). Qualitatively, it appeared that tubulin-GFP fluorescence at the tips of microtubules in the control cells dropped off more slowly to background as compared to the 10 nM Vinblastine-treated cells, in which the fluorescence dropped off more abruptly (Fig. 6B, blue vs yellow). To quantify this observation, we fit a Gaussian error function to the fluorescence intensity drop-off at each microtubule end, as previously described (Coombes et al., 2013; Demchouk et al., 2011), which allowed us to estimate tip tapering that arises as a result of protofilament length variability at the microtubule ends. Consistent with our qualitative observations, we found that the microtubule tip standard deviation was significantly reduced with Vinblastine treatments as low as 1 nM (Fig. 6C; Control vs 1 nM: p=0.004, Z=2.89; Control vs 10 nM: p=6×10^−5^, Z=4.01), indicating that treatment with the microtubule destabilizing drug Vinblastine acted to prune protofilament extensions at growing microtubule plus-ends, eliminating tapered tips with extensive protofilament-edge binding sites. Indeed, tip standard deviations in 10 nM Vinblastine approached the point spread function of the microscope, similar to the blunt ends of GMPCPP microtubules (Fig. 6C (yellow) vs Fig. 1D), suggesting that extended protofilaments and associated tip tapering at growing microtubule ends had been eliminated at higher Vinblastine concentrations.

**Figure 6:**
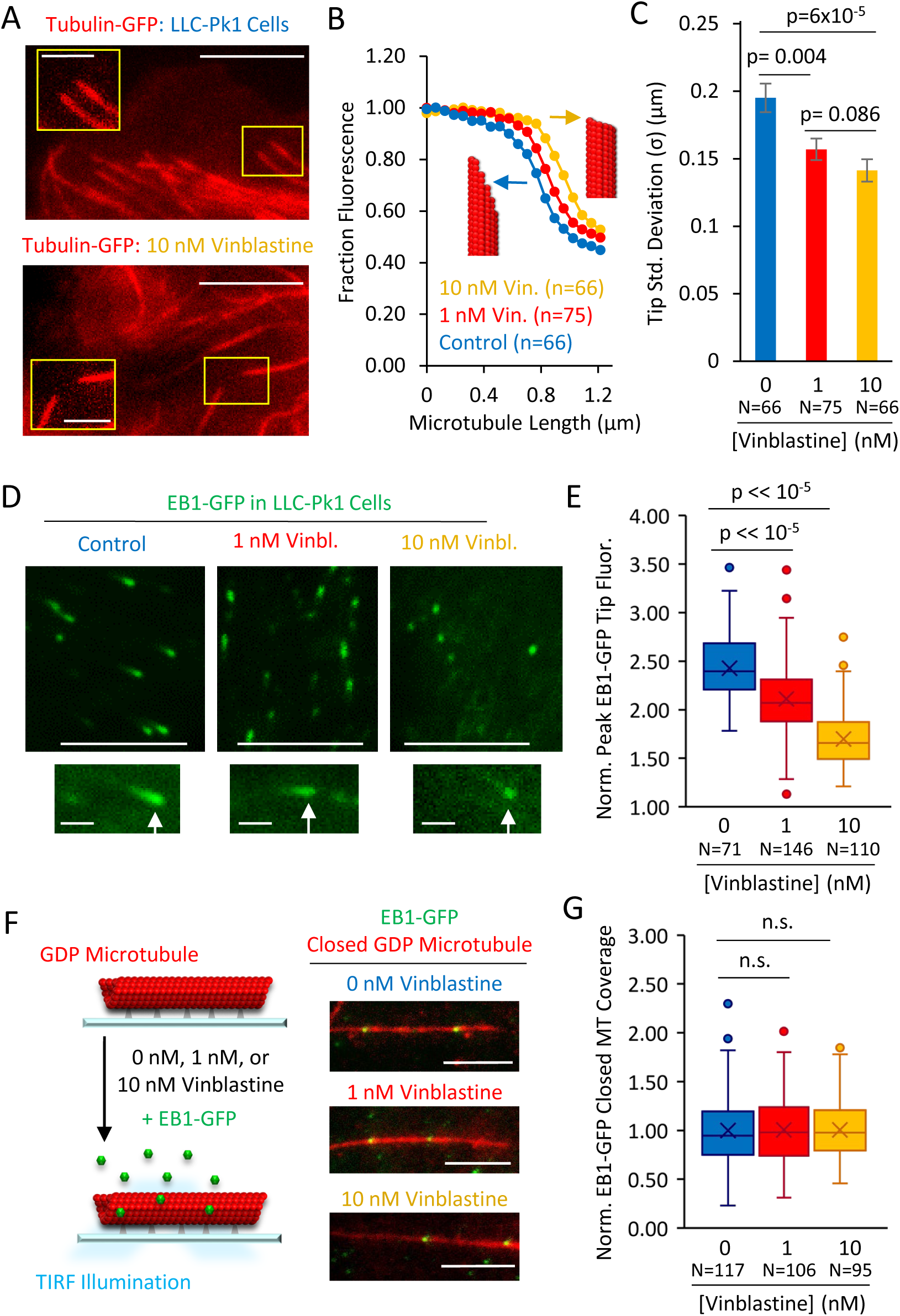
Structural recognition facilitates EB1 tip tracking at growing microtubule plus-ends in cells. (A) Representative images of Tubulin-GFP expressed in LLC-Pk1 cells with inset detail of growing microtubule tips (control cells, top, 10 nM Vinblastine treated, bottom; see Materials and Methods) (scale bar 5 µm, insets 2 µm). (B) Average linescans that show the dropoff in intensity of tubulin at the tips of microtubules in control and Vinblastine treated cells. A sharper drop-off (yellow) is suggestive of a blunt microtubule end, and a slower dropoff (blue), is suggestive of a more tapered microtubule end, as shown by the grey arrows and red microtubule cartoons. (C) Tip standard deviations as estimated by fitting to the graphs in panel F. (D) Representative images of EB1-GFP expressed in LLC-Pk1 cells (top) with inset EB1-GFP comets at growing microtubule tips (bottom) (control cells, left; 1 nM Vinblastine treated, center; 10 nM Vinblastine treated, right; see Materials and Methods) (scale bar 5 µm, comet detail scale bar 1 µm). (E) Peak EB1-GFP fluorescence normalized to lattice EB1-GFP fluorescence. (F) Experiment (left) and representative images (right) of in-vitro experiment to quantify binding of EB1-GFP to closed GDP microtubules in the presence of increasing concentrations of Vinblastine (scale bar, 5 µm). (G) Quantitative binding of EB1-GFP to closed GDP microtubules, based on EB1-GFP binding area on microtubules. All values normalized to the grand average binding area in the 0 nM Vinblastine population. Sample sizes represent number of independent images analyzed (p=0.91, t-test 0 nM vs 1 nM Vin.; p=0.88, t-test 0 nM vs 10 nM Vin.).

We then asked whether blunting of the microtubule tip structure, leading to a reduced number of protofilament-edge binding sites, would alter the efficiency of EB1 tip tracking. Thus, we used an LLC-Pk1 strain with EB1-GFP (Piehl and Cassimeris, 2003), and measured the peak intensity of EB1-GFP at growing microtubule ends, relative to lattice bound EB1-GFP (Fig. 6D; see Materials and Methods). Consistent with qualitative observations (Fig. 6D, bottom, white arrows), we found that peak EB1-GFP intensity at microtubule ends was reduced with increasing Vinblastine concentrations (Fig. 6E, p<<10^−5^, t-test), suggesting that, with flattened end structures, EB1-GFP could not efficiently target to growing microtubule plus-ends. To ensure that the reduced EB1 targeting was not a consequence of shorter EB1-GFP comet lengths in Vinblastine treatment, we then compared the peak intensity of EB1-GFP comets at similar EB1-GFP comet lengths between controls and in 1 nM Vinblastine (Control comet length: 0.60±0.06 µm, n=62; 1 nM Vinblastine comet length: 0.59±0.07 µm, n=90; (mean±SD)). Here, while the EB1-GFP comet lengths were statistically indistinguishable in our samples (p=0.89, t-test), we nevertheless observed a significant decrease in peak comet intensity in the Vinblastine-treated cells relative to the same-comet-length control cells (p=4.5×10^−6^, t-test, Fig. S4A).

Finally, to verify that the reduced peak EB1-GFP comet intensity in Vinblastine was not the result of an overall lower affinity of EB1 for microtubules in the presence of Vinblastine, we measured EB1-GFP binding to closed in-vitro GDP microtubules in the presence of increasing Vinblastine concentrations (Fig. 6F). Importantly, and in contrast to the reduced peak intensity of EB1-GFP comets on dynamic microtubules in cells, the binding of EB1-GFP to closed GDP microtubules was similar regardless of Vinblastine concentration (Fig. 6G, no significant differences, t-test; experiment repeats, Fig. S4B). We conclude that the efficiency of cellular EB1-GFP tip tracking is disrupted with Vinblastine treatment, likely due to blunting of tapered tip structures at growing microtubule ends, which cuts the availability of high-accessibility EB1 protofilament-edge binding sites.

## Discussion

Taken together, our results suggest a new model to explain EB1 tip tracking at growing microtubule plus-ends, in which EB1 recognizes and rapidly binds to the protofilament edges at open, tapered microtubule end structures. The subsequent addition of new GTP-tubulin subunits may then facilitate the maturation of EB1 into a high-affinity binding state on the closed GTP-tubulin lattice. Ultimately, tubulin hydrolysis into a GDP-Pi or GDP state destroys the EB1 binding site, catalyzing its release from the microtubule.

We found that EB1 dwell time distributions on our disrupted structure microtubules were best modeled as two exponential distributions (Fig S2C,D), including one with a short dwell time (∼20 ms), which could be associated with edge-bound EB1. In evaluating this dwell time relative to the kinetics of αβ-tubulin association, in 10 µM tubulin, it has been estimated that a free tubulin subunit would arrive to the growing end of the microtubule as rapidly as every ∼2 ms (Gardner et al., 2011), or as slowly as every ∼10 ms (Mickolajczyk et al., 2019). Therefore, even the relatively short dwell times of edge-bounded EB1 molecules would be long enough to allow for multiple tubulin subunit arrivals while EB1 is bound to a protofilament edge. These kinetics suggest that if EB1 first binds to a protofilament edge, subsequent tubulin subunit arrivals could lock EB1 into its stable 4-tubulin pocket binding site. This pathway to establishing EB1 binding into its 4-tubulin pocket binding location would be efficient both for EB1, in which the steric hindrance for its direct binding into a 4-tubulin pocket was nearly insurmountable in our diffusion simulations, and for incoming tubulin subunits, which would be stabilized longitudinally and/or laterally by binding to EB1 in addition to other protofilament-bound GTP-tubulin subunits.

Overall, we propose that structural recognition by EB1 could promote rapid loading of EB1 at the open, tapered growing microtubule plus-end with its more easily accessible edge sites. Specifically, our results predict that the on-rate for EB1 binding to the closed microtubule lattice would be extremely low, due to very high steric hindrance and low accessibility. In contrast, we predict that the EB1 on-rate would be ∼70-fold (∼6000%) higher at tapered, growing microtubule plus-ends, due to the presence of easily accessible, low steric hindrance sites at exposed protofilament edges. This was the most surprising conclusion from our diffusion simulations: while we expected that tubulin edges would be more accessible to EB1 binding, the enormous magnitude of this preference was surprising to us. Importantly, the binding site geometries and diffusion coefficients used in the simulation were constrained by prior electron microscopy reconstructions and molecule size, and so the molecular diffusion simulations do not rely on any free parameters.

Overall, we propose that structural recognition by EB1 could promote rapid loading of EB1 at the open, tapered growing microtubule plus-end with its more easily accessible edge sites. Specifically, our results predict that the on-rate for EB1 binding to the closed microtubule lattice would be extremely low, due to very high steric hindrance and low accessibility to these closed lattice positions. In contrast, we predict that the EB1 on-rate would be ∼70-fold (∼6000%) higher at tapered, growing microtubule plus-ends, due to the presence of easily accessible, low steric hindrance sites at exposed protofilament edges. After binding to an edge-site, EB1 would likely undergo a higher off-rate as a result of the partial binding interface that is inherent to these edge-sites. However, our results show an increase in steady-state binding to disrupted microtubule structures, regardless of nucleotide state (Fig. 2C-E), suggesting that the overall affinity of EB1 may be higher on disrupted structures, regardless of the likely increase in off-rates. This increase in affinity may be due to a stronger effect of disrupted microtubule structure in accelerating the EB1 on-rate, as compared to increasing the EB1 off-rates. For example, we observed a ∼45-fold increase in EB1 on-rates to disrupted GDP microtubule structures relative to closed microtubule structures (Fig. 3J), while the “short” EB1 dwell times, which could be associated with edge-bound EB1 on GDP microtubules, were only ∼8 fold faster than the lattice-bound “long” dwell times (Fig. S2D), resulting in a net affinity increase of ∼5-fold.

The effect of microtubule structure on microtubule-associated protein binding preference is not likely restricted to EB1, as we find that this preference stems from the physical accessibility of microtubule binding sites, and thus may be pertinent to any polymer binding protein. We expect that the magnitude of this effect will be strongly impacted by the location of the protein binding site on the microtubule lattice, as suggested by the difference in our pocket vs face binding simulations. Here, any protein that has its binding site located in the dip between protofilaments or at the crease between stacked subunits could potentially display microtubule structural recognition. For example, the microtubule-associated protein Doublecortin (DCX) has a very similar binding site to EB1, but is offset by 4 nm vertically (at the α-α-β-β interface). DCX has been reported to recognize microtubule curvature, a configuration that would require an incomplete lattice with exposed protofilament edges to allow for outward protofilament curvature (Bechstedt and Brouhard, 2012; Bechstedt et al., 2014; Fourniol et al., 2010). Similarly, microtubules grown in the presence of CLIP-170 have been associated with the growth of curved oligomers (Arnal et al., 2004), and it has been shown that microtubule lattice defects may be recognized by CLIP-170 to stimulate microtubule rescue (de Forges et al., 2016). Thus, recognition of microtubule curvature or lattice defects by microtubule-associated proteins could rely, at least in part, on an increased arrival rate to exposed protofilament edge sites. In contrast, proteins such as TPX2, which binds in part across the faces of α and β tubulin subunits (Zhang et al., 2017), and TOG1-domain containing proteins such as XMAP215 and Stu2, where the TOG1 domain binds predominantly on the β-tubulin face (Byrnes and Slep, 2017), may be less likely to display structural recognition.

As noted above, because outward protofilament curvature at growing microtubule ends is necessarily associated with exposed protofilament edges, we cannot exclude the possibility that EB1 may also preferentially bind to curved protofilaments (Guesdon et al., 2016). However, we find that decreased steric hindrance can increase the arrival rate of EB1 to protofilament edges by ∼70-fold (∼6000%, Fig. 3), while no similar effect would apply to protofilament curvature. Further, we did not routinely observe outward protofilament curvature at holes and defects within the lattice (Fig. 2B), suggesting that the increased affinity of EB1 to disrupted lattice structures (Fig. 2C) was not a result of outwardly curved protofilaments that were present within the microtubule lattice.

Our simulations model only one microtubule binding domain of EB1, yet EB1 is typically a homodimer. We anticipate that EB1’s dimerization activity would not alter the observed microtubule structural preference in our simulations, but would likely increase the dwell time of edge-associated EB1, as once one domain binds to the more easily accessible protofilament edge sites, EB1’s second binding domain would be constrained to a small area, and therefore much more likely to bind, even within a 4-tubulin pocket, due to the high local availability of binding sites. Thus, the EB1 homodimer as a whole would become more stably bound to the microtubule. EB1 dwell time was not included in our current simulations, as the simulation for a particular molecule was ended upon arrival to the microtubule lattice, but future iterations of our simulation could model dwell time to test this idea.

In conclusion, we find that a high steric hindrance barrier impedes EB1 from binding directly into the pocket-like interface between four adjacent tubulin dimers in the lattice, and that this barrier is markedly reduced at tapered microtubule ends. Protofilament edge-binding greatly increases the arrival rate of EB1 to growing microtubule plus-ends, which facilitates EB1 tip tracking in cells. Finally, our results support a general principle in which microtubule-associated protein binding rates are influenced by the location and conformation of the protein binding interface.

## Acknowledgments

This work was supported by National Institutes of Health grant NIGMS R01-GM103833 and R35-GM126974 and National Science Foundation CAREER award 1350741 to MKG. MZ acknowledges the support of National Institutes of Health grant R35-GM119552, the Human Frontier Science Program and the Searle Scholars Program. The authors have no conflicts of interest to disclose. Parts of this work were carried out in the Characterization Facility, University of Minnesota, a member of the NSF-funded Materials Research Facilities Network (www.mrfn.org) via the MRSEC program. We thank Drs. Joe Howard and Ron Vale for gifting reagents used as part of this project. We thank Holly Goodson and the Gardner Laboratory for helpful discussions.

## Materials and Methods

### Tubulin Purification and Labelling

Tubulin was purified and labelled as per (Gell et al., 2010).

### EB1-GFP Purification

Plasmid pETEMM1-HIS6x-tev-EB1-GFP was transformed into Rosetta(DE3) pLysS E. coli, grown in 10 ml of LB+cam+kan at 37° overnight, and then subcultured into 1L of the same media, mixing at 37° for 2hr to an A600 of 0.44. IPTG was then added to 2 mM and the culture was mixed at 18° for 14 hr. The culture was centrifuged 30 min. at 4°, 4400 xg. The cell pellet was resuspended into 12 ml of PBS / 0.1% tween-20 / 5 mM B-mercaptoethanol / 1 mg/ml lysozyme and protease inhibitors (1 mM AEBSF / 10μM pepstatin A / 0.3μM aprotinin / 10uM E-64), mixed at 4°, 2hr., and then sonicated on ice at 80% power, 50% duty, 10×1 min. The lysate was centrifuged at 4°, 1hr., 18000 xg. The supernatant was passed through 2 ml of Talon Metal Affinity Resin and the resin was sequentially washed with 5 column volumes Buffer A (50 mM sodium phosphate pH7.5 / 300 mM KCl / 10% glycerol / 5 mM B-mercaptoethanol). Next, the column was washed with 95% buffer A/5% bufferB (Buffer A / 300 mM imidazole), followed by a wash with 90% A / 10% B, and finally 85% Buffer A/ 15% Buffer B. All buffers had the protease inhibitors 1 mM AEBSF, 10uM pepstatin A and 10uM E-64). Protein was eluted with 1 ml fractions of 100% Buffer B + above protease inhibitors. Elution fractions were analyzed on coomassie and western blot. Relevant fractions were combined and dialyzed against Buffer A overnight at 4°, then 2hr. at 18° in the presence of HIS6-taggedTEV protease (Expedeon TEV0010) at a 1:100 w/w protein ratio). Dialysate was then mixed with additional Talon resin to remove cleaved HIS6x tag.

### Kif5B-GFP Purification

The RP hk339-GFP construct (monomeric Kif5B-GFP-HIS6x), a gift from Ron Vale (Addgene plasmid #24431, (Tomishige and Vale, 2000), was transformed into the E. coli Rosetta (DE3) pLysS. This strain was cultured in LB+amp+cam at 30° overnight to an A600 of 1.5, then subcultured into 1200 ml of the same media, mixed at 37°, 3hrs to an A600 of 0.47. IPTG was added to 0.2 mM final and growth continued at 16° for 19 hrs. The culture was centrifuged at 2100 xg, 4° for 30 min., supernatant discarded and cells resuspended in lysis buffer (50 mM sodium phosphate pH 6.2 / 250 mM NaCl / 20 mM imidazole / 5 mM B-mercaptoethanol / 1 mM MgCl2 / 0.5 mM ATP / 0.5% triton X-100 plus protease inhibitors (1 mM AEBSF / 10uM pepstatin A / 10uM E-64 / 0.3μM aprotinin / 1 mM benzamidine). Lysozyme was added to 0.1 mg/ml with 150U DNAse I and mixed at 4°, 1hr.The cell suspension was lysed by sonication on ice, 2x 10 min. at 90% power, 50% duty and centrifuged at 4°, 14000 xg for 40 min. The supernatant was pH’ed to 7.6 and mixed with 1 ml Talon Metal affinity resin (Clontech 635509) for 30 min. at 4°. Resin was washed 5x with 2 column volumes of lysis buffer + 0.1x protease inhibitors and protein was eluted with sequential 1 ml volumes of lysis buffer containing 300 mM imidazole pH7 + 0.1x protease inhibitors. The elution fractions were analyzed by coomassie and western blot and the peak fractions used in experiments.

### EB1 conjugation to Gold Beads

Unlabeled EB1 was conjugated to 20 nm gold particles using the Innovacoat Gold Particle labeling Mini Kit (Innovacoat 229-0005) as per kit instructions. EB1 was buffer-exchanged into PBS by centrifuging and washing with a 0.5 ml Amicon Ultra centrifugation filter (30kd cut off). 10μl (5.5μg) of EB1 was diluted with 2μl of kit dilution buffer and 42 μl of kit reaction buffer. 45 μl of this mixture was added to the kit’s gold reagent and allowed to sit 25 min. 5 μl of the kit’s quencher buffer was added and incubated for 5 min. 1 ml of a 1:10 dilution of quencher buffer was added, mixed, and Centrifuged at 4° for 20 min. at 9000 xg. The supernatant was discarded and the gold pellet was suspended in Brb80 buffer.

### Microtubule Pool Preparations

Microtubules for the bound nucleotides GDP, GMPCPP, and GTPγS of both closed and disrupted-structure were prepared as described in (Reid et al., 2017).

Briefly, GDP microtubules were grown using a mixture of 33μM tubulin, 1 mM GTP, 4 mM MgCl_2_, and 4% DMSO, and incubated for 30 min at 37°C. The solution was then diluted 100-fold into BRB80 solution with 10μM Taxol and stored at either 37°C (for closed microtubules) or at 25°C (for disrupted-structure microtubules).

GMCPP microtubules were grown using a mixture of 3.9μM tubulin and 1 mM GMPCPP in BRB80, which was incubated for 1hr at 37°C. To generate disrupted-structure GMPCPP microtubules, closed GMPCPP microtubules were incubated in 40μM CaCl_2_ for 40 minutes immediately before use in an experiment.

GTPγS microtubules were grown using a mixture of 12μM tubulin, 50 mM KCl, 10 mM DTT, 0.1 mg/ml Casein, 4 mM GTPγS, and unlabeled GMPCPP “seed” microtubules to serve as nucleation points. The mixture was incubated for 1hr at 37°C, then diluted 3.5-fold and stored at 37°C overnight. The disrupted-structure GTPγS microtubules were prepared similarly, but with the following changes; the initial mixture contained higher tubulin concentration (25.5μM instead of 12μM), after incubation the mixture was diluted 26-fold instead of 3.5-fold, and was stored at 25°C overnight instead of at 37°C.

### Construction and Preparation of Flow Chambers for TIRF Microscopy Imaging

Imaging flow chambers were constructed as in Section VII of (Gell et al., 2010), with the following modifications: two narrow strips of parafilm replaced double-sided scotch tape as chamber dividers: following placement of the smaller coverslip onto the parafilm strips, the chamber was heated to melt the parafilm and create a seal between the coverslips; typically only three strips of parafilm were used, resulting in two chambers per holder. Chambers were prepared with anti-rhodamine antibody followed by blocking with Pluronic F127, as described in Section VIII of (Gell et al., 2010).

Microtubules were adhered to the chamber coverslip, and the chamber was flushed gently with warm BRB80. The flow chamber was heated to 35°C using an objective heater on the microtubule stage, and then 3-4 channel volumes of imaging buffer were flushed through the chamber.

Microtubules were imaged on a Nikon TiE microscope using 488 nm and 561 nm lasers sent through a Ti-TIRF-PAU for Total Internal Reflectance Flourescence (TIRF) illumination. An Andor iXon3 EM-CCD camera fitted with a 2.5X projection lens was used to capture images with high signal to noise and small pixel size (64 nm). Images were collected using TIRF with a Nikon CFI Apochromat 100X 1.49 NA oil objective.

### EB1-GFP Binding to GMPCPP Microtubules

Closed GMPCPP microtubules, prepared as described above, were introduced into an imaging chamber and allowed 30s-3 min to bind the antibody on the coverslip. The imaging chamber was then flushed with 1X Imaging Buffer. 1X Imaging Buffer consisted of BRB80 supplemented with the following: Casien 80 μg/ml, D-Glucose 20 mM, Glucose oxidase 20 μg/ml, Catalase 10 μg/ml, DTT 10 mM, KCl 30 mM, and Tween-20 1%.

EB1-GFP solution (100 μL Imaging Buffer, 1.5 μL of 11 μM EB1-GFP (163 nM final EB1-GFP concentration) was then introduced into the imaging chamber and allowed 10-15 minutes to bind, after which images were collected in both the microtubule (red) and EB1-GFP (green) channels (561 nm and 488 nm respectively).

### EB1-GFP binding to Microtubule Pools

GDP microtubules, either closed or disrupted-structure, were prepared as described above and introduced into an imaging chamber. Microtubules were allowed 30s-3 min to bind the antibody on the coverslip. The microtubule solution was then flushed out of the chamber with 1X imaging buffer (2X Imaging buffer = 110 mM KCl, 40 μg/ml Glucose Oxidase, 20 μg/ml Catalase, 40 mM D-Glucose, 20 mM DTT, 160 μg/ml Casein, 2% Tween-20, and 20 μM Taxol). EB1-GFP solution (50μL 2X imaging buffer, 45uL BRB80, 5μL of 5μM EB1-GFP) was flowed into the imaging chamber and allowed 10-15 minutes to bind, after which images were collected in both the microtubule (red) and EB1-GFP (green) channels (561 nm and 488 nm respectively).

GMPCPP microtubules, either closed or disrupted-structure, were prepared as described and introduced into an imaging chamber. Microtubules were allowed 30s-3 min to bind the antibody on the coverslip. The microtubule solution was then flushed with low-salt imaging buffer (Low-salt 2X Imaging buffer = 25 mM KCl, 40 μg/ml Glucose Oxidase, 20 μg/ml Catalase, 40 mM D-Glucose, 20 mM DTT, 160 μg/ml Casein, 2% Tween-20, and 20 μM Taxol). EB1-GFP solution (50μL Low-salt 2X imaging buffer, 45uL BRB80, 5μL of 5μM EB1-GFP) was flowed into the imaging chamber and allowed 10-15minutes to bind, after which images were collected in both the microtubule (red) and EB1-GFP (green) channels (561 nm and 488 nm respectively).

GTPγS microtubules, either closed or disrupted-structure, were prepared as described and introduced into an imaging chamber. Microtubules were allowed 30s-3 min to bind the antibody on the coverslip. The imaging chamber was then flushed with 1X imaging buffer (2X Imaging buffer = 110 mM KCl, 40 μg/ml Glucose Oxidase, 20 μg/ml Catalase, 40 mM D-Glucose, 20 mM DTT, 160 μg/ml Casein, 2% Tween-20, and 20 μM Taxol). EB1-GFP solution (50μL 2X imaging buffer, 45uL BRB80, 5μL of 5μM EB1-GFP (250 nM final EB1-GFP concentration)) was flowed into the imaging chamber and allowed 10-15 minutes to bind, after which images were collected in both the microtubule (red) and EB1-GFP (green) channels (561 nm and 488 nm respectively).

### EB1-GFP On-rate and Dwell time movies

Microtubules were prepared as described above and introduced into an imaging chamber. Microtubules were allowed 30s-3 min to bind the antibody on the coverslip. The imaging chamber was then flushed with 1X imaging buffer (2X Imaging buffer = 110 mM KCl, 40 μg/ml Glucose Oxidase, 20 μg/ml Catalase, 40 mM D-Glucose, 20 mM DTT, 160 μg/ml Casein, 2% Tween-20, and 20 μM Taxol). EB1-GFP solution (25μL 2X imaging buffer, 20uL BRB80, 5μL of 5μM EB1-GFP) was flowed into the imaging chamber. Using a capture rate of 100 frames per second, 10 seconds of images were collected per microtubule in both the microtubule (red) and EB1-GFP (green) channels (561 nm and 488 nm respectively).

### Kif5B-GFP Global binding to Microtubules

GDP microtubules, either closed or disrupted-structure, were prepared as described above and then introduced into an imaging chamber. Microtubules were allowed 30s-3 min to bind the antibody on the coverslip. The microtubule solution was then flushed with 1X imaging buffer (2X Imaging buffer = 110 mM KCl, 40 μg/ml Glucose Oxidase, 20 μg/ml Catalase, 40 mM D-Glucose, 20 mM DTT, 160 μg/ml Casein, 2% Tween-20, and 20 μM Taxol). Either 2.5 nM Kib5B-GFP solution (25μL 2X imaging buffer, 14.5uL BRB80, 5μL of 25μM Kif5, and 5μL 100 mM AMPPNP) or 10nM Kif5B-GFP solution (25μL 2X imaging buffer, 14.5uL BRB80, 5μL of 100μM Kif5, and 5μL 100 mM AMPPNP) was flowed into the imaging chamber and allowed 10-15minutes to bind, after which images were collected in both the microtubule (red) and Kif5 (green) channels (561 nm and 488 nm respectively).

### EB1-GFP Binding to GDP and GDP-P_i_ Microtubules

GDP microtubules, either closed or disrupted-structure (prepared as described above), were introduced into an imaging chamber, as described above. The imaging chamber with GDP microtubules was then flushed with either 1X GDP-P_i_ imaging buffer (2X GDP-P_i_ Imaging buffer = 110 mM KPO_4_, 40 μg/ml Glucose Oxidase, 20 μg/ml Catalase, 40 mM D-Glucose, 20 mM DTT, 160 μg/ml Casein, 2% Tween-20, and 20 μM Taxol) or 1X GDP imaging buffer (2X GDP imaging buffer = 110 mM KCl, 40 μg/ml Glucose Oxidase, 20 μg/ml Catalase, 40 mM D-Glucose, 20 mM DTT, 160 μg/ml Casein, 2% Tween-20, and 20 μM Taxol), and were allowed to sit for 20 min. Microtubules were flushed with 1X imaging buffer again, after which an EB1-GFP solution (20 μL 2X GDP-P_i_ or GDP imaging buffer, 13 uL BRB80, 4 μL of 2.5 μM EB1-GFP) was flowed into the imaging chamber and allowed 10 min to bind, after which images were collected in both the microtubule (red) and EB1-GFP (green) channels (561 nm and 488 nm respectively).

### Culture, Imaging, and Drug Treatment of LLC-Pk1 Cells

The LLC-Pk1 cell line expressing EB1-GFP, a gift from Patricia Wadsworth, or expressing GFP-Tubulin, a gift from Lynne Cassemeris, was grown in Optimem media (ThermoFisher #31985070), 10% fetal bovine sera + penicillin/streptomycin at 37° and 5% CO_2_. Cells were grown in 35 mm glass bottom dishes for visualization by microscopy.

For drug treatments, the Optimem media was replaced with CO_2_ independent imaging media and drug dissolved in DMSO, or only DMSO as a control, was added to the final concentration needed. Cells were incubated with the drug/DMSO for 30 min prior to imaging.

### Analysis of EB1-GFP Binding profile on GMPCPP Microtubules

Microtubule and EB1-GFP fluorescence intensity were analyzed, and the microtubule aligned, as previously described (Coombes et al., 2016). Briefly, the single time point images of GMPCPP microtubules with EB1-GFP were cropped to separate each microtubule into a single image using ImageJ. Then, integrated and averaged line scans of each microtubule image were created using a MATLAB script. In each case, the microtubule was aligned with the brighter EB1-GFP (green) fluorescence end on the right, and then the red and green fluorescence were plotted as a function of microtubule length from the dimmer EB1-GFP signal end to the brighter EB1-GFP signal end. To do this, the green EB1-GFP fluorescence was integrated along the length of each microtubule ± 256 nm above and below the microtubule centerline to account for point spread function and variability in properly finding the microtubule centerline. Then, the green fluorescence intensity was summed over the last 576 nm on both ends of the microtubule. The lower summed value was considered the dimmer EB1-GFP end, while the higher summed value was deemed the brighter EB1-GFP end. To combine all individual microtubule data into an ensemble average plot, the microtubules were rebinned to a common length, represented by the mean length of all observed microtubules (Gardner et al., 2005). Scatter plots of the ensemble average values were created by importing the integrated line scan fluorescence data into Excel.

### Microtubule Tip Structure Analysis

Average tip standard deviation values were calculated by fitting an error function to the microtubule ends, as previously described (Demchouk et al., 2011).

### EB1-GFP/Tubulin Binding Ratio Analysis

The average EB1-GFP/Tubulin binding ratio was analyzed using a custom MATLAB script. Briefly, the ends of each microtubule image were manually clicked by a user. Then, the average green and red fluorescence intensity between the clicks, including +/- 4 pixels above and below the centerline (256 nm) established by the two clicks, was calculated. Background and noise in the red and green channels was estimated by calculating the average green and red fluorescence between the two clicks, but in the pixels 8-22 above the centerline established by the two clicks, and in the pixels 8-22 below the centerline established by the two clicks. For each microtubule, the average EB1-GFP signal over background was calculated by dividing the average signal along the microtubule by the average background estimate. Similarly, the average red microtubule signal over background was calculated by dividing the average red microtubule signal by the average red background estimate. Finally, the binding ratio of average green signal over background to average red signal over background was calculated and reported for each microtubule.

### EB1-GFP Binding Coverage Analysis

To compare relative binding of EB1-GFP on GDP and GDP-P_i_ microtubules, and in the presence of Vinblastine, the total length of green (EB1-GFP) occupancy was divided by the total length of the red microtubules on each image (defined as EB1-GFP “microtubule coverage”). This was accomplished by using previously described semi-automated MATLAB analysis code (Reid et al., 2017). Briefly, first, automatic processing of the red microtubule channel was used to determine the microtubule-positive regions, which then allowed for conversion of the red channel into a binary image with white microtubules and a black background. The green EB1-GFP channel was then also pre-processed to smooth high-frequency noise and to correct for TIRF illumination inhomogeneity. The green channel threshold was then manually adjusted to ensure visualization of all EB1-GFP binding areas on each microtubule. Measurements of the total EB1-GDP length were then automatically collected from the identified microtubule regions. For presentation, the EB1-GFP microtubule coverage values for each image and condition were normalized to a specific grand average control value.

### Analysis of Peak EB1-GFP Binding to Microtubules in LLC-Pk1 Cells

To analyze the peak EB1-GFP binding to growing microtubule ends, individual comets were analyzed using MATLAB. EB1-GFP comet movies were first analyzed to ensure that only mature comets were selected for analysis. For each mature comet, MATLAB code was used which allowed two manual clicks on each comet: at the location of the highest intensity spot on each comet, and on the microtubule lattice just outside the end of the comet. In addition, the user clicked twice to measure the length of the comet. Then, the fluorescence intensity at the highest intensity spot on the comet was divided by the fluorescence intensity on the microtubule lattice outside of the comet, which we defined as the normalized peak EB1-GFP tip fluorescence.

### Electron Microscopy Experiments

To collect electron microscopy images of GMPCPP microtubules, the microtubules were prepared as described above. One drop of the microtubule solution was placed on a 300-mesh carbon coated copper grid for a duration of 60 seconds, after which the grid was stained with 1% uranyl acetate for 60 seconds. The stain was the wicked away using filter paper, and the grid was left to dry and then stored until use. The samples were imaged using an FEI Technai Spirit BioTWIN transmission electron microscope. Cryo-EM was performed on the same electron microscope. Samples were prepared on a 300-mesh copper grid with a lacey-carbon support film. Grids were treated in a Pelco Glow Discharger before the addition of the GDP microtubule sample and freezing in vitreous ice using a FEI Vitrobot.

Electron microscopy imaging of EB1 conjugated to gold beads was performed by placing one drop of a solution containing disrupted-structure GDP microtubules (prepared as described above) with gold-bead-conjugated EB1 onto a 300-mesh carbon coated copper grid. The sample grid was treated identically to the procedure described above for TEM of GMPCPP microtubules.

### Analysis of EB1-bead experiment

Images of EB1 conjugated to gold beads were analyzed in two parts. First, each image was examined manually. Gold beads appearing in an image were tallied and classified according to their proximity to a microtubule and the appearance of the microtubule at the location of the bead. Only beads directly contacting a microtubule were tallied, and were classified into the following categories: Edges, for gold beads localized at the edges of sheet-like regions of a microtubule or at a microtubule end; Defect, for gold beads localized to regions of the microtubule with visible defects such as gaps or breaks; and Lattice, for gold beads localized to intact regions of the microtubule. Additionally, gold beads found on sheet-like regions were sub-classified based on their location on the sheet, as either Sheet-Edge or Sheet-Middle. To be classified as a Sheet-Edge, a bead was required to overlap with the sheet edge but have its center located beyond, not overlapping, the sheet. Secondly, using a custom MATLAB script, the total perimeter of the all microtubules in the gathered images was traced and each segment was classified as being Lattice or Edge/Defect, based on how a bead would have been classified if it was located at the given segment of microtubule. The lengths for each classification were summed and used to normalize the counts.

### EB1 3D Diffusion and Binding Simulation: Setup and Assumptions

Approximations for the shapes of a tubulin dimer and EB1 microtubule binding domain (PDB ID: 3JAR (Zhang et al., 2015)) were created manually based off of Cryo-EM reconstruction data using 3D modeling software (Blender) and exported to an .obj object file format. EB1’s binding site coordinates were determined by calculating the center of the contacting faces for each of the four adjacent tubulin dimers, and the binding center was determined as the average position of the four binding interface coordinates.

The simulation was coded and run using MATLAB 2015a (Mathworks). The tubulin dimer subunits were arranged into the canonical microtubule arrangement, with either a blunt or tapered end, maintaining 207 tubulin dimers (average protofilament length ∼16 dimers) for both conditions. EB1 was initialized at a random position on a sphere of radius 500 nm centered on the microtubule. We note that while the simulated microtubule shape was approximately cylindrical with a height of ∼128 nm, the random initial localization of EB1 within a 500 nm radius of the microtubule did not bias the simulated EB1 molecules towards the microtubule ends, as demonstrated by our face-binding simulations (Fig. 4F), in which no end bias was observed.

### EB1 3D Diffusion and Binding Simulation: Dynamic calculation of time step size

In order to run the simulation efficiently, the time step for the simulation was dynamically adjusted based on the distance of EB1 from the microtubule, similar to (Castle and Odde, 2013). Briefly, the time step was large (∼7×10^−6^ sec) when the simulated EB1 molecule was far from the microtubule (≥100 nm), and small (∼ 4×10^−12^ sec) when the EB1 molecule was close to the microtubule (≤ 1 nm). Specifically, the scaled time step sizes were calculated based on the translational diffusion speed of the EB1 molecule, where the translational diffusion step size was governed by:

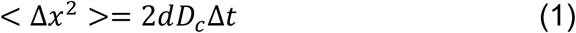

Where *Δx*=translational diffusion molecule step size, *d* = dimensions (=3), *D_c_*=translational diffusion coefficient (see below), and *Δt*=time step size. This equation was then used to determine the scaled time step size, such that when the EB1 molecule was ≥100 nm from the microtubule (eg, *Dist_i_* ≥ 100), the time step size was then calculated from Eq. 3 such that the molecule would be 5 steps away from the microtubule (eg, *Δx_i_ ≡ Dist_i_ / 5*), and when EB1 was ≤ 1 nm from the microtubule, the time step size was then calculated from Eq. 3 such that the molecule would be 20 steps away from the microtubule (eg, *Δx_i_ ≡ Dist_i_ / 20*). The time step size was scaled linearly between those lengths, and so that a step closer to the microtubule reduced the time step size, until the EB1 molecule was 1 nm from a binding site, at which time a fixed time step was used that resulted in typical diffusion lengths of 0.5 angstroms per step. The distance used in these calculations (*Dist_i_*) was the separation distance between the closest points on the microtubule and on EB1, not the center-to-center distance.

### EB1 3D Diffusion and Binding Simulation: EB1 3D Diffusion

Each EB1 molecule was allowed to diffuse both translationally and rotationally until the molecule was either further than 2000 nm from the simulation center, or else had the EB1 binding interface properly oriented and within 1 nm of the binding interface center of any microtubule binding location. EB1 diffusion was governed by the calculated translational and rotational diffusion coefficients as determined by equations (2) and (3) respectively:

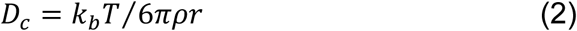

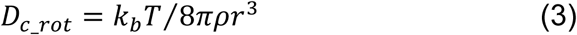

In the above equations, ρ is viscosity of water, and r is the radius of the EB1 molecule, approximated as the average radius of all its vertex points (2.2 nm). Diffusion and rotation was assumed to be equal across all axes.

The translational distance that the EB1 molecule traveled in a time step was then calculated based on random sampling from a Gaussian distribution governed by:

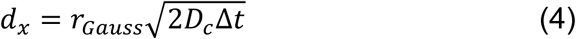

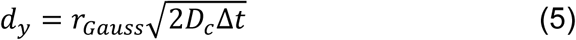

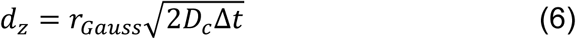

Where d_x_, d_y_, and d_z_ were the translational displacement in x, y and z respectively, and r_Gauss_ was a normally distributed random number (mean: 0, standard deviation: 1).

Similarly, in the same time step, the angle of rotation for EB1 was also determined from normally distributed random number, and the rotational diffusion constant:

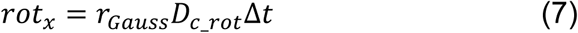

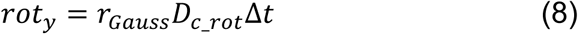

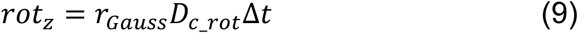

### EB1 3D Diffusion and Binding Simulation: EB1 Collision with a Microtubule

EB1 was not permitted to pass through or intersect volumes with any tubulin dimers. The dynamic time steps prevent skipping from one side of a tubulin dimer to the other in one time step, thus avoiding collision, by maintaining appropriately small step sizes when near a tubulin dimer. Collisions and measurements of the nearest point distance between EB1 and any tubulin subunit were conducted using the Voronoi-Clip (vClip) algorithm (Mirtich, 1998).

Briefly, the closest point between the EB1 polyhedron and a tubulin dimer polyhedron were determined by randomly selecting initial candidate features (vertex, edge or face) for each object. Then, the neighboring features of the EB1 candidate were tested to find one that was closer to the Tubulin candidate feature. If a feature was found to be closer than the initial candidate, it became the new candidate. This was repeated for both the EB1 and Tubulin objects until no two candidates could be found that were closer together than the current pair. Then, if the distance between them was less than or equal to zero (with respect to the object’s outer surface), that indicated that they were intersecting and a collision had occurred, in which case the EB1 position was reverted to the previously recorded value, and the EB1 molecule permitted another attempt at diffusion without collision. If the distance between candidate features was greater than zero, no collision had occurred and the simulation proceeded to the next time step.

The vClip method is useful in reducing computational load, because the final candidate features can be stored from the last time step and used as the initial candidate features for the next test (Mirtich, 1998). Because the time steps are small, the saved candidate features are often the optimal candidate features (or adjacent to the optimal candidate features) for the new position (that has moved and rotated only slightly).

A second optimization technique was used to avoid testing against every tubulin dimer at every time step. Briefly, a much more rapid test for center to center distance was calculated for EB1 to detect “nearby” tubulin dimers (determined by the sum of the maximum vertex radius for both EB1 and the tubulin dimer, and a buffer distance) as compared to the more expensive and accurate vClip test. “Nearby” was determined by a “Sweep and Prune” algorithm, specifically a hierarchical binary tree of tubulin dimer bounding-boxes that was used to very quickly determine if any tubulin dimers needed be considered “Nearby”.

### EB1 3D Diffusion and Binding Simulation: EB1 Binding to a Microtubule

EB1 stopped diffusing, and the simulation was ended for that molecule, when its binding interface was *oriented properly* and within 1 nm of a binding interface on the microtubule. This “binding” configuration was constrained by both distance and molecule orientation relative to its binding site on the microtubule: if the binding interface was oriented *away* from the microtubule, the separation distance could be as little as 0 nm, and the binding interface would still be >4 nm (EB1 diameter) away from the tubulin pocket binding interface. In this situation, the simulation would not stop, but the EB1 molecule would continue to diffuse and rotate until it entered into the proper binding position, or diffused out of the simulated volume.

---

**Figure S1: Supplemental data to support Figure 1.**
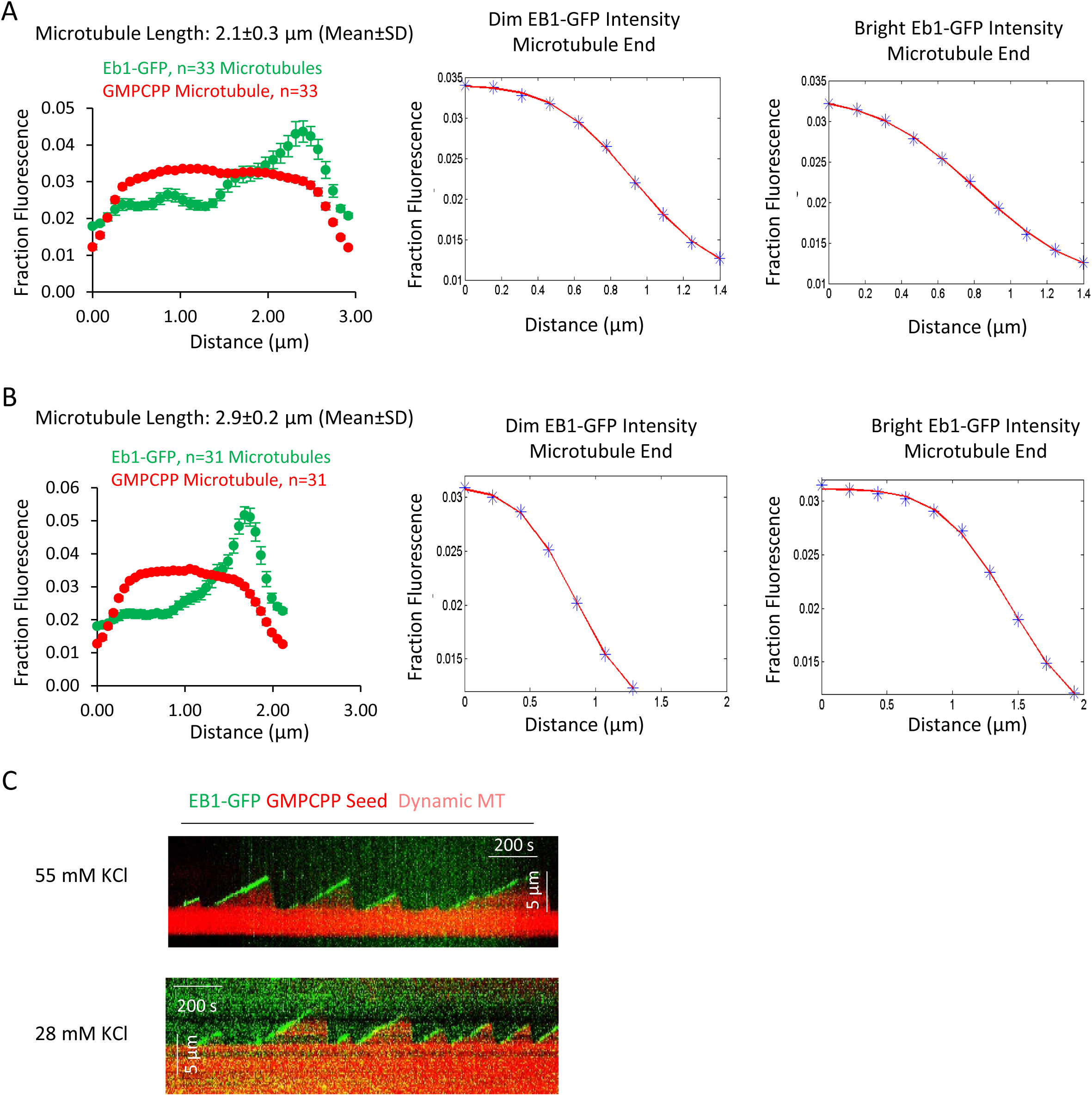
(A and B) Averaged fluorescence line scan data for EB1-GFP and rhodamine-labeled GMPCPP microtubules of length 2.1 ±0.3 μm (A) and 2.9 ±0.2 μm (B). The microtubule end with brighter green fluorescence intensity was aligned on the right side of the graph, with the dimmer green intensity on the left. Center: Ensemble-averaged microtubule intensity data (blue points), and fitted curve (red line) for the dimmer EB1-GFP microtubule end. Right: Ensemble-averaged microtubule intensity data (blue points), and fitted curve (red line) for the brighter EB1-GFP microtubule end. Note that the end with brighter EB1-GFP binding had a slower (more lengthy) transition from full intensity to background intensity in the red (microtubule) channel, indicating a more extended transition from full lattice to extended protofilaments, as compared to the dimmer EB1-GFP end. Fits were determined using MATLAB and were fitted to an error function. (C) Example kymographs demonstrating EB1-GFP tip tracking of dynamic microtubules, both at 55 mM KCl, and at 28 mM KCl.

**Figure S2: Supplemental Data to support Figure 3:**
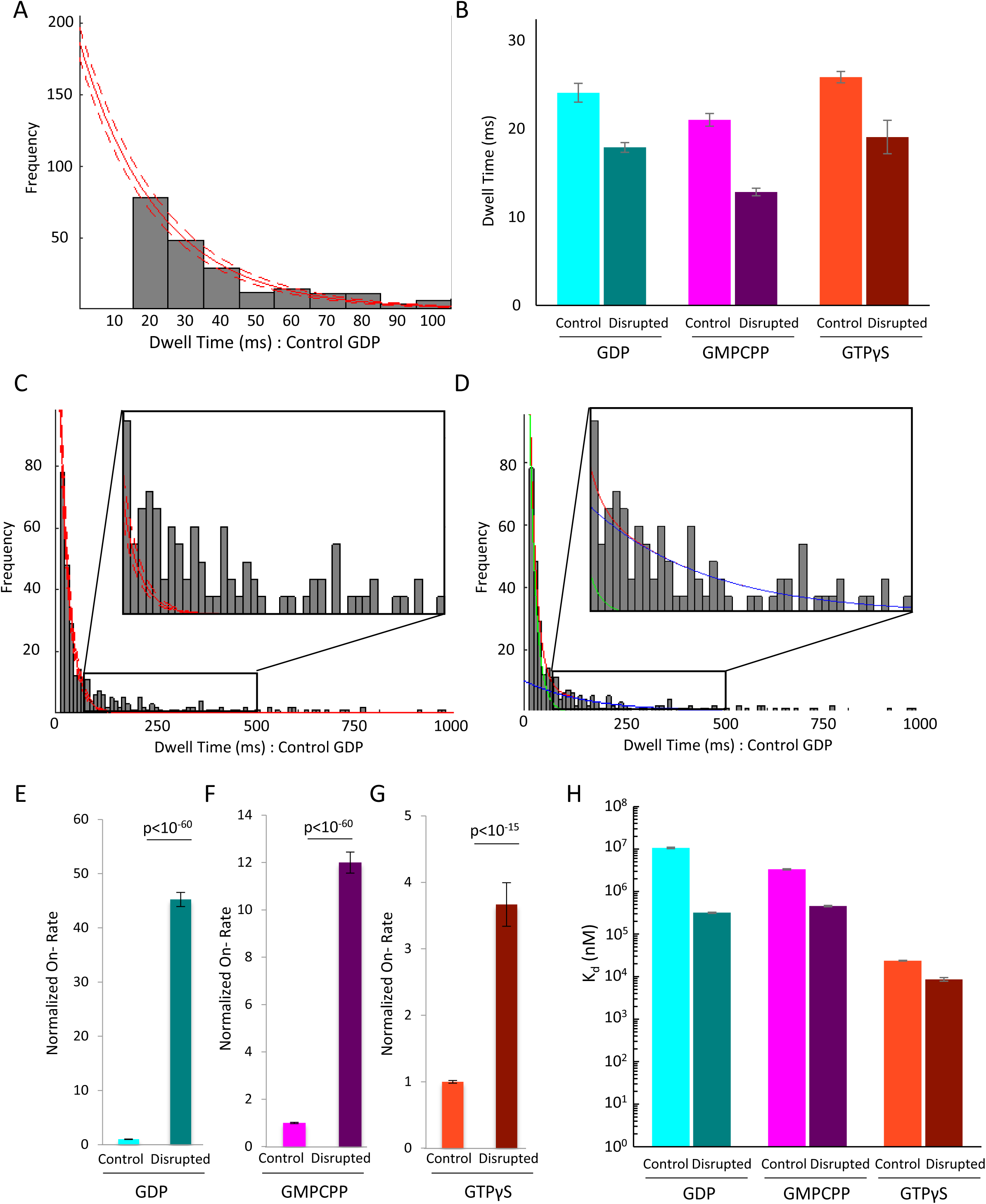
(A) Clipped histogram of EB1-GFP dwell times on control GDP microtubules (longer dwell time bins clipped off). EB1-GFP was observed with a frame-rate of 10 ms. Single-frame binding events (dwell time ≤ 10 ms) were not included in fitting due to higher likelihood of missed events and false positives (unfilled bar at t=10 ms). A single exponential fit (solid red line) and associated 95% confidence intervals (dashed red lines) were used to account for the uncounted, short binding events. Histograms for other nucleotides and structural conditions were fit in the same way (not shown). (B) Characteristic EB1-GFP dwell times as determined from the exponential fit, shown for control and disrupted pools in GDP (cyan, teal), GMPCPP (magenta, purple) and GTPγS (orange, brown) microtubules. Error-bars show the 95% confidence interval for the dwell-time fit parameter. In all cases, the disrupted-structure microtubules had a slight but significant decrease in the dwell-times for EB1-GFP as compared to the control microtubules, as would be expected with an increase in the proportion of binding sites with fewer than the full complement of four tubulin binding partners at edges. All microtubule types had characteristic dwell times on the order of a 10-25 ms, which is in agreement with previous findings that EB1 has a very short dwell time at microtubule ends. (C) Full histogram of control GDP microtubules as shown in panel A, but including the longer-dwell time bins. Inset shows dwell time histogram from 80 ms to 500 ms. At this scale it becomes apparent that the single exponential fit (the fitting of which did include the longer dwell time data points) does not reproduce to frequency of dwell times >100 ms. (D) Same histogram as in panel C, but fitted with the sum of two exponential curves. The first curve (green line) maintains a characteristic dwell time that is the approximately the same as the initial fitting from panel A, ∼15 ms. The second curve (blue line) has a much longer characteristic dwell time (∼150 ms) and a much smaller Y intercept. The sum of these two curves (red line) very closely matches the observed distribution and does reproduce the subpopulation of longer dwell-time binding events. This finding matches well with our hypothesis and model that EB1’s low dwell time is due largely to binding events at microtubule edges, which have fewer binding partners and are thus more likely to rapidly dissociate, whereas binding events at full lattice sites are proportionally less common, but would be more stably bound when they do occur. (E-G) Normalized on-rate values for EB1-GFP for the six different microtubule conditions. In each case, the data is normalized to the mean control population on-rate value. Error bars are SEM, derived from errors of the coefficients from exponential fit of dwell-times. (H) Calculated dissociation constant (K_d_) values for the six pools of microtubules. Despite the very similar dwell times for all six pools, the differences in on-rate leads to K_d_ values that decrease from GDP to GMPCPP to GTPγS microtubules, and with each disrupted pool having a lower K_d_ than in the control pool.

**Figure S3: Supplemental Data to support Figure 5:**
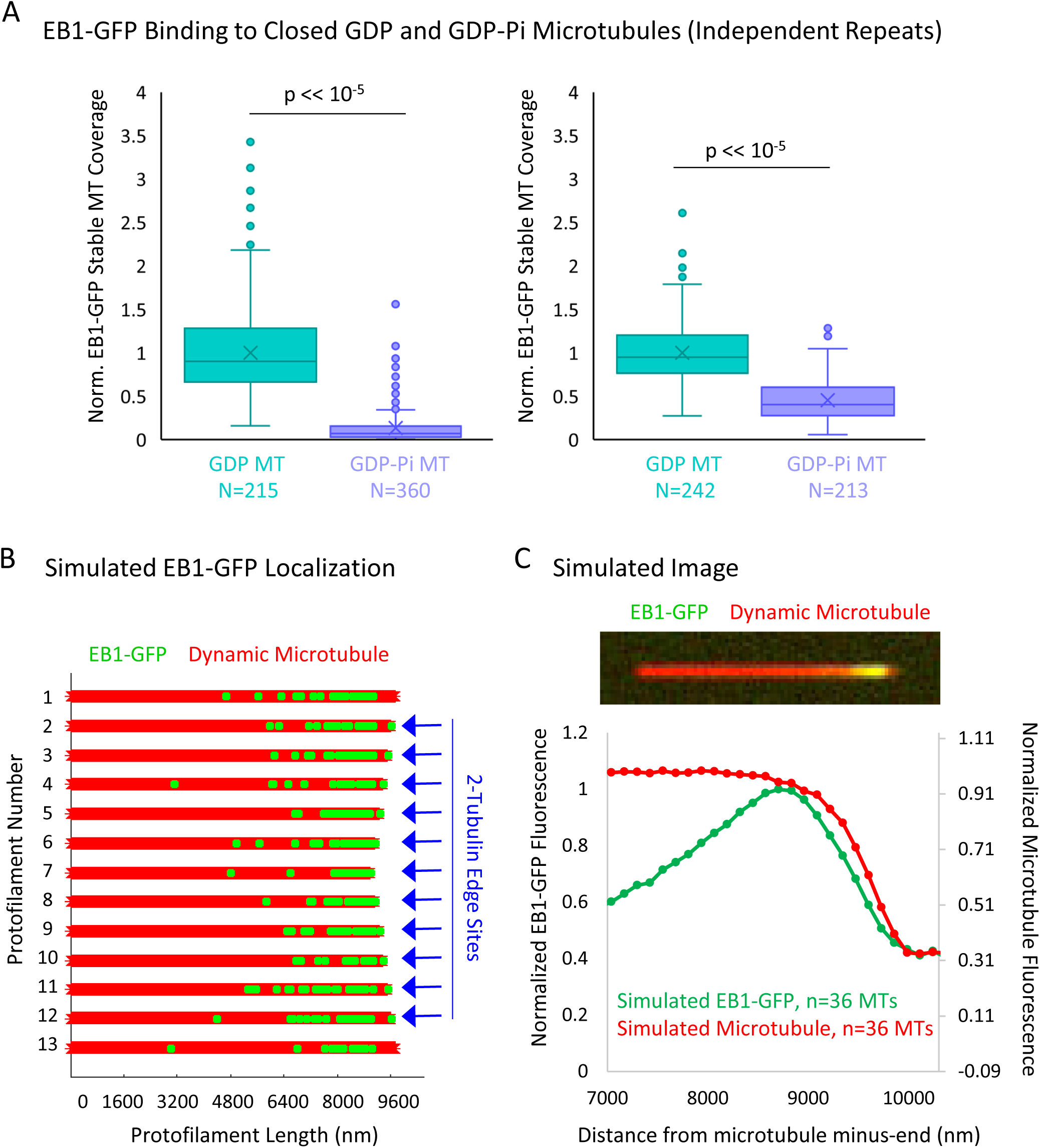
(A) Experimental repeats to evaluate the binding of EB1 to closed GDP and GDP-Pi microtubules. Each plot represents a different experimental day. (B) Graphical representation of a flattened microtubule, showing microtubule protofilaments (red) with EB1-GFP at the microtubule plus-end (green). The microtubule shown has moderate taper at its plus-end (at the right of schematic). EB1-GFP is localized at the microtubule plus-end via two rules: (1) the EB1-GFP is distribution decreases exponentially away from the tip, starting at the first complete layer of tubulin subunits, and (2) the 2-tubulin “protofilament edge” sites are fully occupied by EB1-GFP. Locations with 2-tubulin edge sites are noted with blue arrows. The longest two protofilaments (1 and 13) are not assigned edge sites to account for a closed tube. (C) Top: Simulated image of a microtubule (red) with EB1-GFP at its end (green). Bottom: Super-averaged line scan data from 36 simulated microtubules with varying degrees of tip taper. Similar to previous reports (Maurer et al., 2014), the peak in EB1-GFP appears to be slightly penultimate to the microtubule end, suggesting that binding to protofilament edges may not significantly bias the observed EB1-GFP fluorescence distribution at the microtubule plus-end.

**Figure S4: Supplemental Data to support Figure 6:**
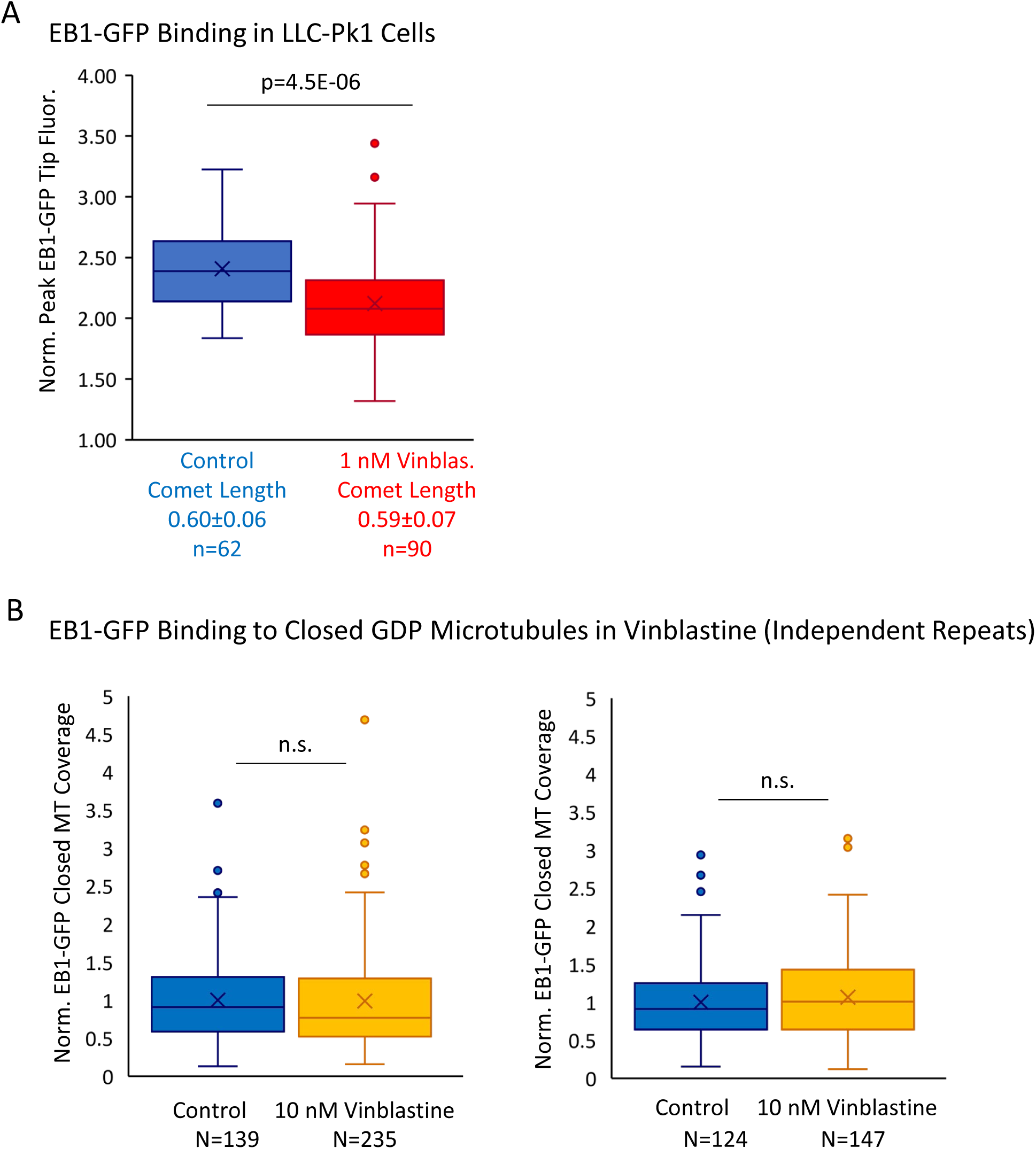
(A) Peak EB1-GFP fluorescence normalized to lattice EB1-GFP fluorescence for a subselection of LLC-Pk1 cellular EB1-GFP comets with similar lengths in control and 1 nM Vinblastine. (B) Experimental repeats to evaluate the binding of EB1 to closed GDP microtubules in the presence of 0 nM (control) and 10 nM Vinblastine. Each plot represents a different experimental day.

